# Enhanced aversive memory retrieval by chemogenetic activation of locus coeruleus norepinephrine neurons

**DOI:** 10.1101/831313

**Authors:** Ryoji Fukabori, Yoshio Iguchi, Shigeki Kato, Kazumi Takahashi, Satoshi Eifuku, Shingo Tsuji, Akihiro Hazama, Motokazu Uchigashima, Masahiko Watanabe, Hiroshi Mizuma, Yilong Cui, Hirotaka Onoe, Keigo Hikishima, Yasunobu Yasoshima, Makoto Osanai, Ryo Inagaki, Kohji Fukunaga, Takuma Nishijo, Toshihiko Momiyama, Richard Benton, Kazuto Kobayashi

## Abstract

The ability to retrieve memory store in response to the environment is essential for animal behavioral adaptation. Norepinephrine (NE)-containing neurons in the brain play a key role in the modulation of synaptic plasticity underlying various processes of memory formation. However, the role of the central NE system in memory retrieval remains unclear. In this study, we developed a neural chemogenetic activation strategy using insect olfactory Ionotropic Receptors (IRs), and used it for selective stimulation of NE neurons in the locus coeruleus (LC) in transgenic mice. Ligand-induced activation of LC NE neurons resulted in enhancement of the retrieval process of conditioned taste aversion, which was mediated through at least partly adrenergic receptors in the amygdala. Pharmacological blockade of LC activity confirmed the facilitative role of these neurons in memory retrieval. Our findings indicate that the LC-amygdalar pathway is required and sufficient for enhancing the recall of taste associative memory.

## Introduction

The ability to retrieve necessary information associated with environmental stimuli and context is indispensable for animal behavioral adaptation. Disturbances in this information retrieval process, especially in humans, result in degradation of not only quality of life but also the sense of personal identity (***Klein and Nichols, 2012***).

Impairments specific to the retrieval process have been reported in some amnesiac patients, who exhibit the inability to explicitly recall test stimuli despite their spared implicit memory of the same stimuli, which can be retrieved with prompts or partial information (***Warrington and Weiskrantz, 1970***). By contrast, recurrent involuntary memory retrieval after traumatic events is one of the major symptoms of post-traumatic stress disorder (***American Psychiatric Association, 2013***). A number of clinical and preclinical studies have suggested that multiple neurotransmitter/modulator systems are implicated in the retrieval process (***Kopelman, 1992***), but the detailed neural mechanism of this process is not well understood.

Norepinephrine (NE)-containing neurons in the brain are divided into discrete cell groups in the pons and medulla, projecting to a diverse array of brain regions (***Chandler et al., 2014; Robertson et al., 2013***). NE plays a key role in the modulation of long-lasting synaptic potentiation and strengthening in the hippocampus (***Gelinas and Nguyen, 2005; Huang and Kandel, 1996; O’Dell et al., 2010***) and amygdala (***Huang et al., 2000; Huang and Kandel, 2007; Johansen et al., 2014***). Indeed, a number of behavioral studies demonstrate that the central NE system contributes to associative aversive memory processes, including acquisition (***Bahar et al., 2003; Bush et al., 2010; Ferry et al., 2015***), consolidation (***Guzmán-Ramos et al., 2012; LaLumiere et al., 2013***), and reconsolidation (***Kobayashi et al., 2000; Villain et al., 2016; Zhou et al., 2015***).

By contrast, studies of the NE system in the retrieval process with different behavioral tasks have produced controversial results. Electrical stimulation of the locus coeruleus (LC), the major NE cell group in the brain (***Chandler et al., 2014; Robertson et al., 2013***), enhances performance in a complex maze task during a retention test, which is blocked by administration of a β-adrenergic receptor antagonist (***Devauges and Sara, 1991; Sara and Devauges, 1988***). A genetic study with knockout mice for dopamine β-hydroxylase showed that NE is involved in the retrieval of a particular type of contextual and spatial memory dependent on the hippocampus (***Murchison et al., 2004***). However, application of a β-adrenergic receptor antagonist into the amygdala does not influence the retrieval of conditioned flavour aversion requiring amygdalar function (***Miranda et al., 2007***). Therefore, the exact role of the central NE system in memory retrieval still remains unclear.

In this study we address the role of NE neurons, focusing on the LC in the retrieval process of conditioned taste aversion. We developed a novel chemogenetic approach to activate specific neuronal types that employs olfactory Ionotropic Receptors (IRs) from *Drosophila melanogaster* (***Abuin et al., 2011; Grosjean et al., 2011***). This approach, which we have named INTENS (<u>in</u>sect ionotropic r<u>e</u>ceptor-mediated <u>n</u>euronal <u>s</u>timulation), enables efficient and sustained stimulation of the target neurons. The INTENS technology achieved the activation of LC NE neurons expressing IRs in response to exogenous ligands, and the resultant stimulation of NE release enhanced retrieval of conditioned memory for taste. This role of the LC NE neurons in memory retrieval was mediated at least in part through α_1_- and β-adrenergic receptors in the amygdala. Pharmacological experiments that inhibit LC NE activity confirmed the facilitative role of these neurons in memory retrieval through the LC-amygdalar pathway via adrenergic receptor subtypes.

## Results

### Insect IR-mediated stimulation of LC NE neurons

IRs constitute a family of sensory ligand-gated ion channels distantly related to ionotropic glutamate receptors and mediate environmental chemical detection in *Drosophila melanogaster* and other insects (***Benton et al., 2009; Rytz et al., 2013***). The heteromeric complex composed of IR84a and IR8a subunits confers excitatory cellular responsiveness to odorants such as phenylacetaldehyde (PhAl) and phenylacetic acid (PhAc) (***Abuin et al., 2011; Grosjean et al., 2011***). We used this odorant-specific reaction of the IR84a/IR8a complex for the INTENS technology (***Figure 1A***).

**Figure 1.**
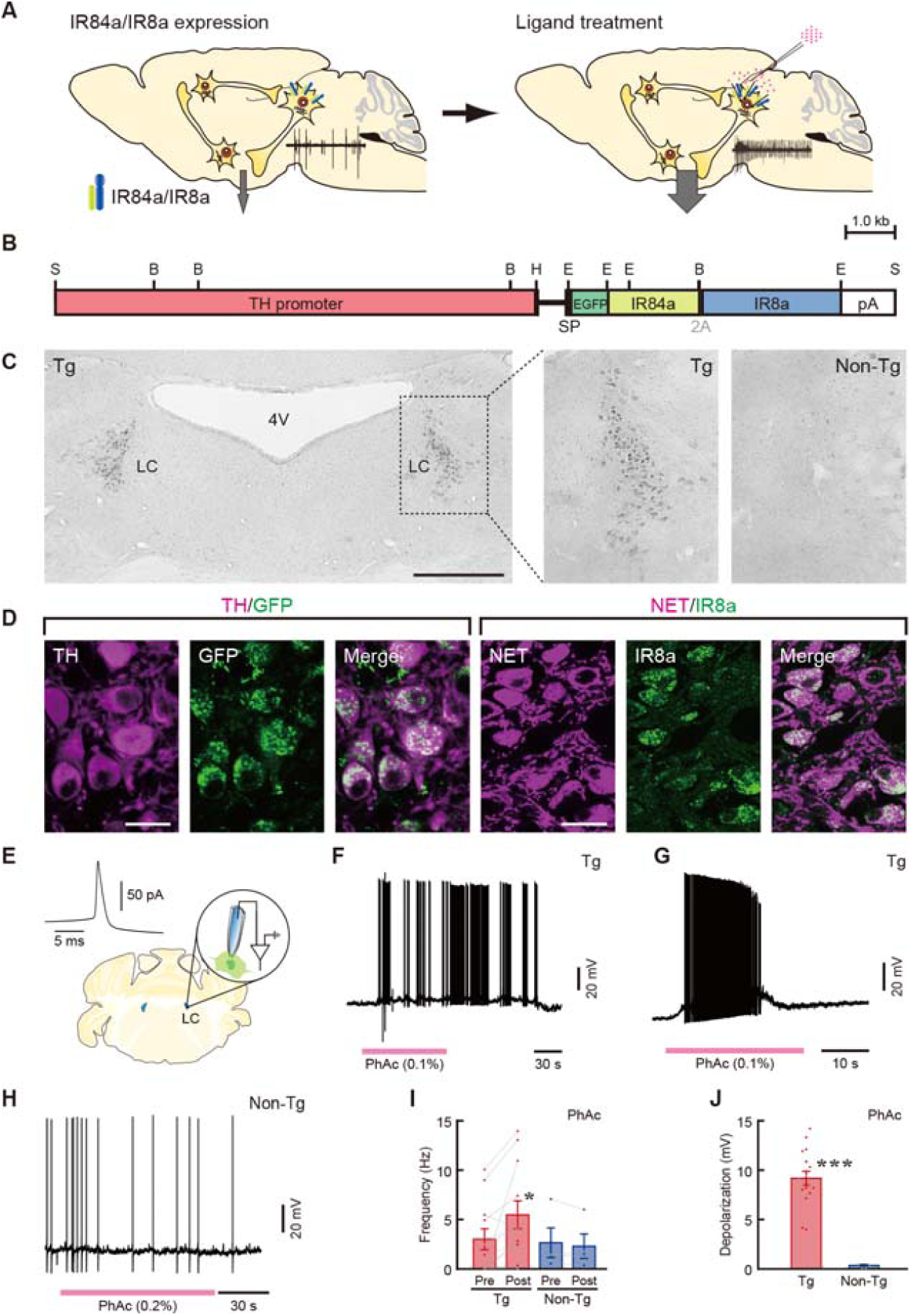
Experimental strategy, transgene expression, and ligand-induced LC activation in Tg mice. (**A**) Strategy for INTENS. Tg animals expressing IR84a/IR8a genes in specific cell types are treated with exogenous ligands into the target brain regions. (**B**) Structure of the gene cassette encoding EGFP-IR84a-2A-IR8a with a signal peptide (SP) downstream of the TH gene promoter. B, *Bam*HI, E, *Eco*RI, H, *Hin*dIII, S, *Sal*I. pA, polyadenylation signal. (**C**) Transgene expression in the LC revealed by GFP immunohistochemistry. 4V: fourth ventricle. (**D**) Confocal microscopic images of LC sections obtained from double immunohistochemistry for TH/GFP and NET/IR8a. Scale bars: 500 μm (C), 20 μm (D). (**E**) Schematic illustration of strategy for a whole-cell current-clamp recording of a brain slice preparation. Inset shows a typical waveform of NE neurons showing a wide action potential with large after-hyperpolarization. (**F**) Excitatory effect of PhAc (0.1%) on the membrane potential of a NE neuron obtained from a Tg mouse. (**G**) Depolarization block after excitatory response to PhAc in another neuron in a Tg mouse. (**H**) Lack of effect of PhAc (0.2%) on the membrane potential of a neuron from a non-Tg mouse. (**I, J**) Bar graphs showing the firing frequency before (pre) and after (post) PhAc application (**I**) and amplitude of PhAc-induced depolarization (**J**) of NE neurons in Tg (n = 12 for firing frequency and n = 17 for depolarization amplitude) and non-Tg (n = 4) mice. *p < 0.05 vs. pretreatment in Tg mice (paired two-tailed t-test), ***p < 0.001 vs. non-Tg mice (unpaired two-tailed t-test). Data are presented as mean ± SEM. Individual data points are overlaid.

IR84a/IR8a genes are expressed under the control of a cell type-specific gene promoter, and the target neurons are stimulated with exogenous ligands (PhAl/PhAc). To express the receptors in NE neurons, we generated transgenic (Tg) mice carrying the genes in which IR84a fused to enhanced green fluorescent protein (EGFP) was connected to IR8a via a 2A peptide, downstream of the tyrosine hydroxylase (TH) gene promoter (***Sawamoto et al., 2001****; **Matsushita et al., 2002***) (***Figure 1B***). Immunostaining for GFP indicated the transgene expression in LC neurons in the TH-EGFP-IR84a/IR8a mouse line, whereas there was no expression in the non-Tg littermates (***Figure 1C***, ***Figure 1 — figure supplement 1A and B*** for expression patterns in other catecholamine-containing cell groups). In the Tg mice, EGFP-IR84a expression was colocalized with TH immunoreactivity, and IR8a reactivity was colocalized with signals for NE transporter (NET) (***Figure 1D***). Double immunostaining for GFP and TH showed that almost all LC neurons express these receptors (93.6 ± 0.02% TH^+^/GFP^+^ cells/total TH^+^ cells, n = 4). *In situ* hybridization with antisense probes also confirmed both IR84a and IR8a gene expression in the Tg-mouse LC (***Figure 1 — figure supplement 1C***).

To examine cellular responsiveness of LC NE neurons expressing IR84a/IR8a, we performed a whole-cell current-clamp recording in slice preparations (***Figure 1E***). NE cells in the LC were identified by a low firing frequency (< 7 Hz) and wide action potentials with large after-hyperpolarization, as observed in previous studies (***van den Pol et al., 2002; Zhang et al., 2010***). In *Drosophila*, robust activation of IR84a/IR8a-expressing cells was induced by treatment with 0.1% (8.3 mM) PhAl (***Abuin et al., 2011***). We used 0.1% solution of the ligands for bath application in slices. Bath application of 0.1% PhAl depolarized the membrane and increased the firing frequency of NE neurons in the LC of the Tg mice, whereas it had little or no effect on the membrane potential of the non-Tg LC neurons (see ***Figure 1 - figure supplement 2***).

The application of 0.1% (7.3 mM) PhAc also depolarized the membrane of LC NE neurons in Tg mice, increasing the firing frequency (***Figure 1F***). In 5 out of 17 neurons examined, the firing stopped after depolarization had reached its steady state even in the presence of PhAc (***Figure 1G***), indicating the occurrence of depolarization block in these neurons. In non-Tg mice, PhAc had little or no effect on the firing frequency or membrane potential of LC neurons (***Figure 1H***). Excluding the neurons showing depolarization block, the firing frequency in Tg mice was significantly elevated from 3.04 ± 1.04 Hz (pre) to 5.47 ± 1.38 Hz (post) by the PhAc application (***Figure 1I***, n = 12, paired two-tailed t-test, t_11_ = 2.667, p = 0.0219). In non-Tg mice, the frequency was similar between pre- and post-PhAc application (2.63 ± 1.48 Hz and 2.35 ± 1.25 Hz, respectively) (***Figure 1I***, n = 4, paired two-tailed t-test, t_3_ = 1.011, p = 0.3864). The amplitude of PhAc-induced depolarization was 9.25 ± 0.69 mV including the neurons in which depolarization block was observed, and significantly greater compared to the non-Tg value (0.34 ± 0.08 mV) (***Figure 1J***, Levene’s test, F_(1, 20)_ = 5.060, p = 0.0365, unpaired two-tailed t-test with Welch’s method, t_16.42_ = 12.92, p < 0.0001). The baseline firing of LC neurons in the preparations used for PhAc application appeared to show higher frequency than in the case of PhAl, but the average of these values ranged within ∼7 Hz as the definition of LC NE neurons, and the values were also indistinguishable between the Tg and non-Tg groups in each preparation. The data obtained from the *in vitro* electrophysiology indicated that application of exogenous ligands induced excitatory cellular responsiveness of LC NE neurons expressing IR84a/IR8a. The INTENS technique enables us to stimulate the firing activity of specific neuronal types by treating with these ligands.

To confirm this ligand-specific reaction of the receptors in mammalian cultured cells, we generated a lentiviral vector to express IR84a/IR8a complex. HEK293T cells were transduced with the lentiviral vector, and the expression of the two receptors was detected by immunohistochemistry (***Figure 1 — figure supplement 3A***). A whole-cell voltage-clamp experiment indicated that PhAc application induced a dose-dependent increase in the inward currents in the cells (***Figure 1 — figure supplements 3B and C***). In addition to the slice electrophysiology, the data support the suggestion that IR84a/IR8a complex forms a functional receptor in response to exogenous ligands in the mammalian system.

### *In vivo* activation of LC neurons stimulates NE release

To explore whether the INTENS technique can trigger *in vivo* noradrenergic activation, we performed an extracellular single-unit recording of LC NE neurons (***Figure 2A***). NE cells in the LC were identified based on firing frequency and waveform as described (***Takahashi et al., 2010***). For the *in vivo* electrophysiology, we used the concentration of 1% of PhAc for pneumatic injection. The baseline firing rates (pre) of the Tg and non-Tg control mice were not significantly different from each other (1.13 ± 0.24 Hz and 1.20 ± 0.21 Hz, respectively; unpaired two-tailed t-test, t_9_ = 0.205, p = 0.8425). The injection of PhAc into the LC induced a robust increase of firing activity in Tg mice, whereas LC neurons in non-Tg controls were insensitive to PhAc injection (***Figure 2B*** for typical firing patterns). The firing rate of Tg neurons was significantly increased to 3.73 ± 0.31 Hz (post) by the PhAc injection (***Figure 2C***; n = 6, paired two-tailed t-test, t_5_ = 11.53, p < 0.0001 vs. pre), whereas the rate of non-Tg mice did not alter by the injection (***Figure 2C***; post: 1.22 ± 0.15 Hz; n = 5, paired two-tailed t-test, t_4_ = 0.300, p = 0.7788 vs. pre). When the effect of PhAc on the discharge of non-NE neurons around the LC was tested, these neurons were categorized into two groups showing low (0.1 to 4 Hz) and high (9 to 40 Hz) frequency of firing activity. The firing rate in each group was unaffected by PhAc injection (***Figure 2D***, n = 5 or 7, paired two-tailed t-test, t_4_ = 0.460, p = 0.6696 for low frequency, t_6_ = 0.880, p = 0.4125 for high frequency). These data indicated that ligand treatment indeed activates the *in vivo* firing of LC NE neurons harbouring IR84a/IR8a.

**Figure 2.**
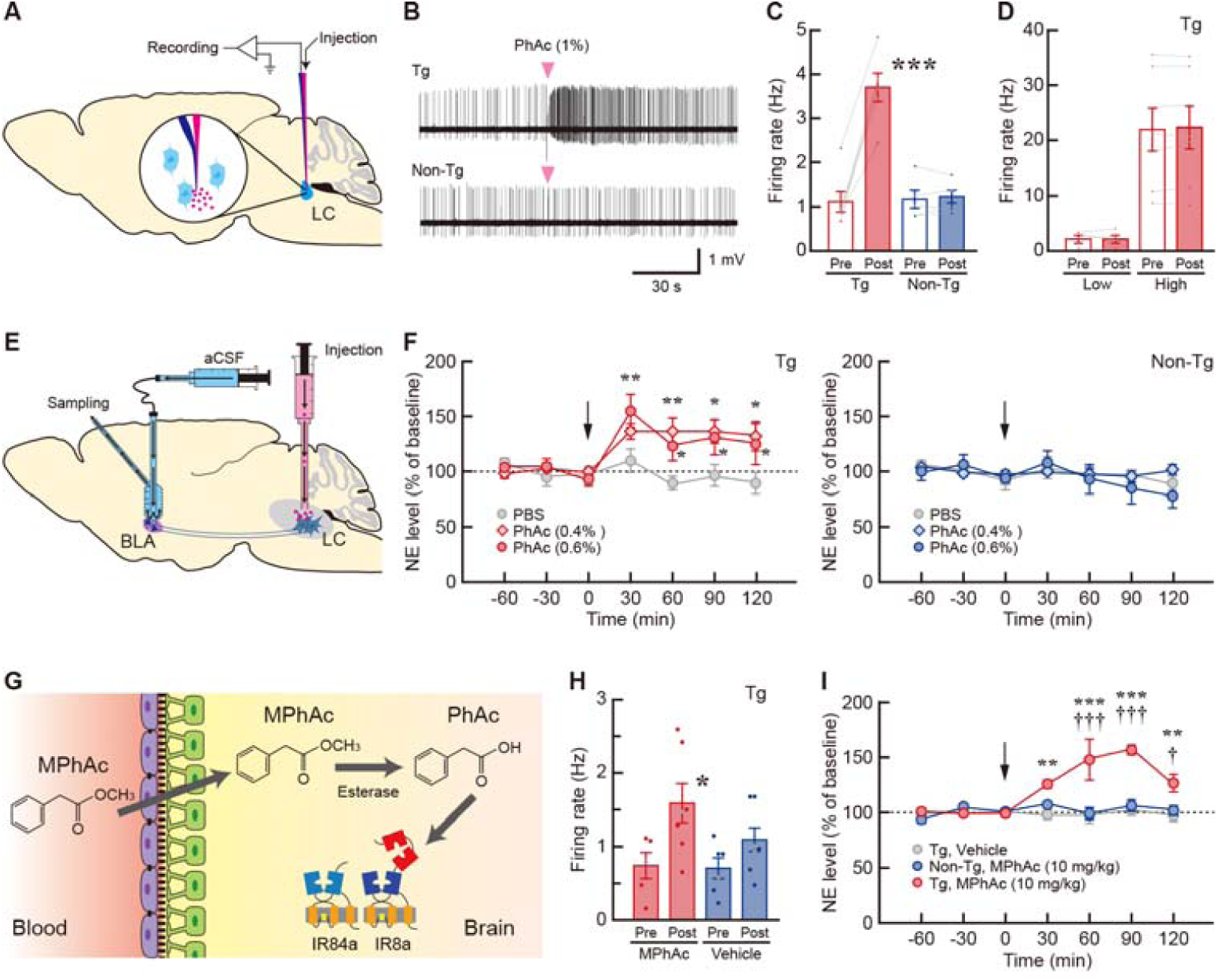
*In vivo* LC activation and increased NE release in Tg mice. (**A**) Schematic diagram for the strategy of single-unit recording. PhAc solution was pneumatically injected through a glass capillary attached to the recording electrode. (**B**) Firing pattern of an identified NE neuron in the Tg (upper panel) and non-Tg (lower panel) mice. Timing of PhAc injection is indicated. (**C**) Firing rate in the pre- and post-PhAc injection period for the Tg (n = 6) and non-Tg (n = 5) mice. ***p < 0.001 (paired two-tailed t-test). (**D**) Firing rate of non-NE neurons in the Tg mice with low and high frequency of the activity in the pre- and post-PhAc injection period (n = 5 or 7 for each type). (**E**) Diagram for the microdialysis for measuring NE release in the BLA in response to LC activation. PhAc solution (0.4/0.6%) or PBS was injected into the LC, and dialysis samples were collected from the BLA. (**F**) Changes in the extracellular NE level after PhAc injection in the Tg and non-Tg mice. NE levels are expressed as a percentage of each animal’s average baseline levels. n = 5 or 6 for each group. *p < 0.05, **p < 0.01 vs PBS (Holm-Bonferroni test). (**G**) Model of systemic drug delivery. MPhAc is systemically administered and transferred across the blood-brain barrier. MPhAc is metabolized in the brain by esterase activity into PhAc, which stimulates IR84a/8a complex. (**H**) Firing activity of LC neurons in the Tg mice in the pre- and post-periods for systemic administration of MPhAc (n = 5 or 7 for each group) or vehicle (n = 6 or 7 for each group). *p < 0.05 vs pretreatment of MPhAc-administered mice (unpaired two-tailed t-test). (**I**) Extracellular NE level after systemic injection of MPhAc or vehicle. NE levels expressed as a percentage of each animal’s average baseline levels. n = 4 for each group. **p < 0.01, ***p < 0.001 vs vehicle-treated Tg mice, ^†^p < 0.05, ^†††^p < 0.001 vs MPhAc-treated non Tg (Holm-Bonferroni test). Data are presented as mean ± SEM.

To further assess ligand-induced noradrenergic activation, we measured NE release in the brain region innervated by the LC using a microdialysis procedure. We first tested NE release from the anterior cingulate cortex (ACC) (***Figure 2 - figure supplement 1A***), because the ACC is known to receive abundant innervations from the LC (***Chandler et al., 2014; Robertson et al., 2013***). When we performed LC microinjection with 0.1% PhAc in this microdialysis with Tg mice, the extracellular NE level did not show a significant increase in the ACC (***Figure 2 - figure supplement 1B***). We then used higher concentrations of PhAc (0.4% and 0.6%) for the microinjections. LC stimulation with these concentrations of PhAc induced a remarkable increase of extracellular NE levels from the baseline in the ACC of Tg mice (***Figure 2 - figure supplement 1C***). Therefore, we used 0.4/0.6% PhAc for LC microinjection and measured NE release from the basolateral nucleus of the amygdala (BLA) (***Figure 2E***). There was no significant difference in average tonic NE concentration (pg/sample) between the Tg and non-Tg mice during baseline fractions before the microinjection: Tg, 1.14 ± 0.32 (n = 16), non-Tg, 1.61 ± 0.18 (n = 15) (unpaired two-tailed t-test, t_29_ = 1.229, p = 0.2288). Injection of both 0.4% and 0.6% PhAc into the LC caused a rapid and long-lasting increase in the extracellular NE level in the Tg mice (***Figure 2F***, n = 5 or 6 for each group, two-way mixed-design ANOVA, drug effect: F_(2, 13)_ = 5.160, p = 0.0224, fraction effect: F_(6, 78)_ = 6.598, p < 0.0001, interaction: F_(12, 78)_ = 2.318, p = 0.0137). The NE level at the 30-min fraction was significantly increased to approximately 155% of the baseline level for 0.6% PhAc injection (Holm-Bonferroni test, t_91_ *=* 3.216, p = 0.0054 vs PBS). The 0.4% PhAc injection also increased the NE level at the 30-min fraction by approximately 136%, although this effect did not reach the statistical significance (t_91_ = 1.871, p = 0.0645 vs PBS). Both 0.6% and 0.4% PhAc induced a sustained elevation in the NE level following the fraction immediately after injection, for 60-min (t_91_ = 2.495, p = 0.0144 for 0.6% vs PBS, t_91_ = 3.420, p = 0.0028 for 0.4% vs PBS), 90-min (t_91_ = 2.478, p = 0.0150 for 0.6% vs PBS, t_91_ = 2.819, p = 0.0177 for 0.4% vs PBS), and 120-min fractions (t_91_ = 2.529, p = 0.0132 for 0.6% vs PBS, t_91_ = 3.055, p = 0.0089 for 0.4% vs PBS). In the non-Tg animals, both 0.6% and 0.4% PhAc injections did not generate any significant changes in NE release (***Figure 2F***, n = 5 for each group, two-way mixed-design ANOVA, drug effect: F_(2, 12)_ = 0.583, p = 0.5733, fraction effect: F_(6, 72)_ = 1.743, p = 0.1234, interaction: F_(12, 72)_ = 0.621, p = 0.8180). One week after the microdialysis experiments, the animals were subjected to histological analysis to validate specific/non-specific toxicity of PhAc treatment to the mouse brain. Immunostaining for TH/NET and cresyl violet staining did not indicate any non-specific damage to LC neurons treated with 0.4%/0.6% PhAc (***Figure 2 - figure supplement 2A–C***). Staining of cell death markers also confirmed the lack of cytotoxicity of PhAc treatment (***Figure 2 - figure supplement 2D***). These data confirmed LC noradrenergic activation by the INTENS strategy, which resulted in the stimulated NE release in the nerve terminal regions innervated by the LC.

Additionally, we tested whether exogenous ligands could stimulate specific neuronal types expressing the receptors in the brain through systemic administration. Since methyl ester derivatives are known to possess high permeability through the blood-brain barrier (***Shukuri et al., 2011; Suzuki et al., 2004; Takashima-Hirano et al., 2010***), methyl phenylacetic acid (MPhAc) was used as a ligand precursor, which is converted to PhAc by esterase activity in the brain (***Figure 2G***).

MPhAc solution (20 mg/kg) or vehicle was intravenously administered in the Tg mice and then used for *in vivo* electrophysiology (***Figure 2H***). The firing rate of LC neurons was significantly elevated from 0.74 ± 0.18 Hz (pre) to 1.59 ± 0.27 Hz (post) by the MPhAc administration (n = 5-7, unpaired two-tailed t-test, t_10_ = 2.376, p = 0.0389), whereas there was no significant difference in the rate between the pre- and post-vehicle administration (0.72 ± 0.14 Hz and 1.08 ± 0.17 Hz, respectively; n = 6-7, unpaired two-tailed t-test, t_12_ = 1.589, p = 0.1380). Next, microdialysis samples collected from the ACC were analysed for NE release. Systemic administration of MPhAc resulted in a marked increase in the extracellular NE level in the Tg mice (***Figure 2I*,** n = 4 for each group, two-way ANOVA, group effect: F_(2, 9)_ = 24.06, p < 0.0001, fraction effect: F_(6, 54)_ = 6.807, p < 0.0001, interaction F_(12, 54)_ = 5.584, p < 0.0001), and NE level at each fraction was significantly higher than the level of vehicle-administered Tg mice (30-120 min, ts_63_ > 3.385, ps < 0.05) or MPhAc-administered non-Tg (60-120 min, ts_63_ > 2.894, ps < 0.05). These data suggest that systemic administration of a ligand precursor, through translocation across the blood-brain barrier, efficiently stimulates NE activity in the Tg animals.

### Ligand-induced LC activation results in enhanced memory retrieval

To address the role of LC NE neurons in memory retrieval, we conducted selective activation of these neurons using INTENS and investigated its impact on the retrieval process of conditioned taste aversion, in which animals learn an association between a taste stimulus and a visceral malaise-inducing stimulus (***Bermúdez-Rattoni, 2004; Yamamoto et al., 1994***). We used a taste reactivity test, which is a sensitive marker of taste aversion (***Inui et al.,2013; Yasoshima and Shimura, 2017***) (***Figure 3A*** for the behavioral procedure and ***Figure 3B*** for the experimental apparatus). In this paradigm, taste aversion memory was formed by repeated conditioning, in which the voluntary consumption of 0.5 M sucrose as a conditioned stimulus (CS) was followed by intraperitoneal 0.15 M lithium chloride (LiCl) injection (2% of body weight) as an unconditioned stimulus (US). After the habituation period, animals received bilateral injections of PBS or PhAc solution into the LC 20 min prior to the intraoral infusion of the CS fluid. We used 0.4% and 0.6% PhAc, which induced a significant increase in NE level in the microdialysis experiment. Animal behavior was recorded, and the latency to express the rejection responses (gaping, chin rubbing, forelimb flailing, paw wiping, and CS dropping) was measured. The latency to express the responses in the Tg mice differed significantly among the treatments into the LC (***Figure 3C***, n = 8-9 for each group, F_(2, 23)_ = 7.214, p = 0.0037), and the values for the 0.4% PhAc (95.25 ± 14.25 s, p = 0.0189) and 0.6% PhAc (79.22 ± 12.39 s, p = 0.0050) injections showed a significant shortening compared to the PBS injection (196.78 ± 35.91 s). The latency for the PhAc injection into the non-Tg LC at the same concentrations (187.88 ± 42.62 s for 0.4% PhAc and 180.25 ± 35.88 s for 0.6% PhAc) demonstrated no significant differences compared to the PBS injection (189.09 ± 36.58 s) (***Figure 3C***: n = 8-11 for each group, F_(2, 24)_ = 0.0149, p = 0.9853). These results suggest that PhAc-induced activation of LC NE neurons enhances the retrieval process of conditioned taste aversion.

**Figure 3.**
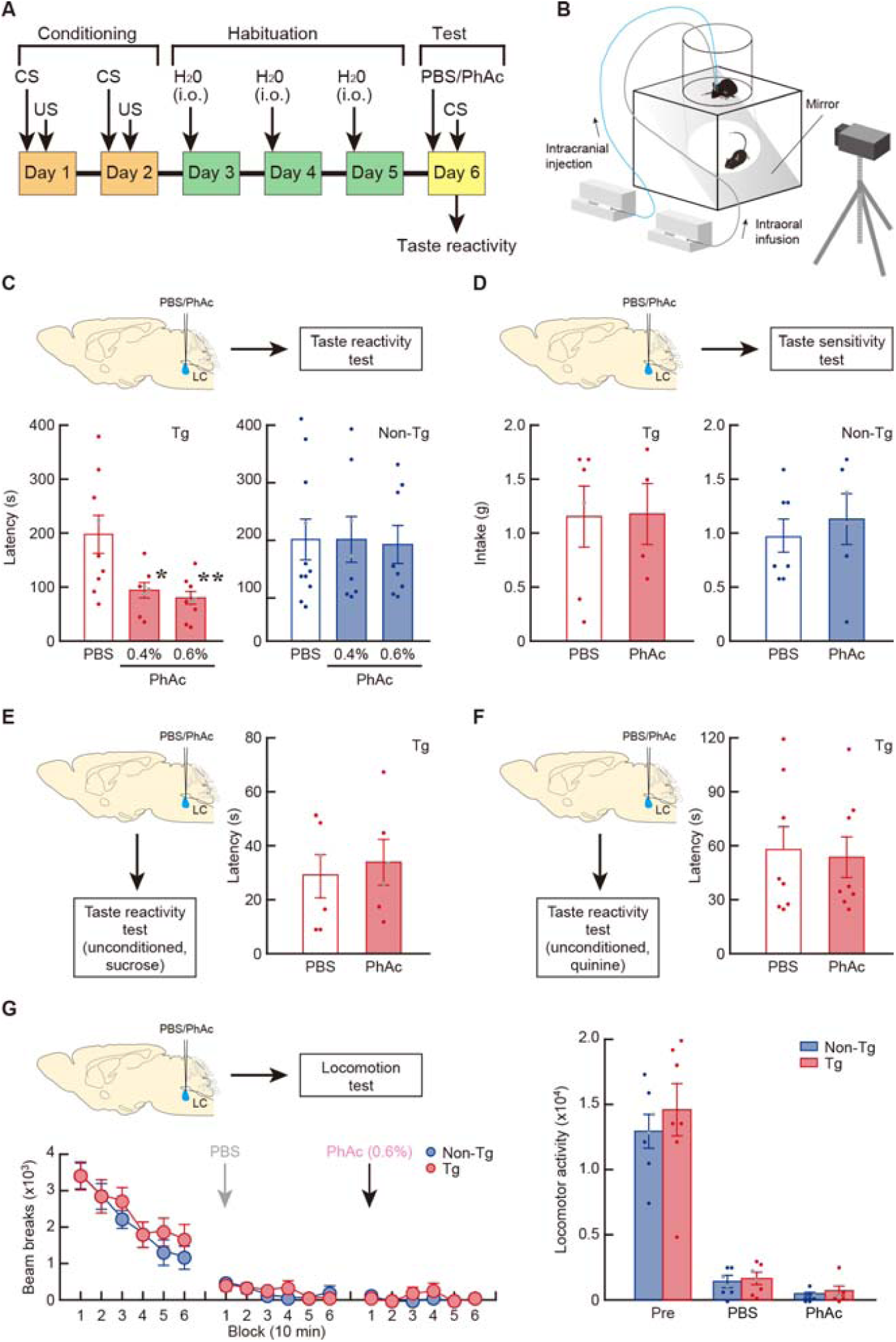
Ligand-induced LC activation enhances taste memory retrieval. (**A)** Schedule of the taste reactivity test. In the conditioning phase, mice were presented 0.5 M sucrose as the CS, followed by an intraperitoneal injection with 0.15 M LiCl as the US. Then, the mice were habituated to intraoral infusion with tap water in the test chamber. During the test phase, mice received a bilateral LC injection of PBS or solution containing PhAc (0.4/0.6%) followed by CS presentation to evaluate the rejection response. (**B**) Experimental apparatus used for the taste reactivity test. A mouse was placed in the test chamber and infused intraorally through a syringe pump, and animal’s behavior was monitored from the bottom through an inside mirror using a digital video camera. (**C**) Taste reactivity test showing shorter latency of rejection response by ligand-induced LC activation in Tg mice. n = 8 or 9 for each group in Tg mice. n = 8-11 for each group in non-Tg mice. *p < 0.05, **p < 0.01 vs PBS in Tg mice (Tukey HSD test). (**D**) Taste sensitivity test presenting normal intake of 0.5 M sucrose in unconditioned mice. PBS or PhAc solution (0.6%) was bilaterally injected into the LC, and the fluid intake of 0.5 M sucrose was measured. n = 4-7 for each group. (**E**) Taste reactivity test showing normal hedonic responses to 0.5 M sucrose in unconditioned Tg mice. PBS or PhAc solution (0.6%) was bilaterally injected into the LC, and the latency for hedonic responses was measured. n = 6 for each group. (**F**) Taste reactivity test displaying unaltered aversive responses to 0.2 mM quinine in unconditioned Tg mice. PBS or PhAc solution (0.6%) was injected into the LC, and the latency for rejection responses was measured. n = 8 for each group. (**G**) Locomotor activity. The number of beam breaks was counted for every 10-min session. The total number of beam breaks during a 60-min test period was calculated as locomotor activity during the pretreatment (Pre) and after PBS treatment (blocks 1-6) and the following 0.6% PhAc treatment (blocks 7-12). n = 7 for each group. Data are presented as mean ± SEM. Individual data points are overlaid.

During the recording of animal behavior, we counted the number of rejection responses. The time course of the responses during the test period indicated that the responses in the Tg mice were generated at earlier phases after 0.4% or 0.6% PhAc injection compared to the PBS injection, although the total number of aversive responses during the test period was similar in the three treatment groups (***Figure 3 — figure supplement 1***). The data suggest no apparent change in memory storage by the LC stimulation in our experimental condition. Therefore, in the following analyses the effect of ligand-induced LC activation on memory retrieval was evaluated by measuring the latency for rejection behaviors.

The reduction in the latency to express the rejection responses may be attributable to increased sensitivity to taste stimulus. To test this possibility, we examined whether the activation of LC NE neurons changes the sensitivity of taste by measuring the consumption of 0.5 M sucrose presented to unconditioned animals. Mice received the bilateral injection of PBS or PhAc solution (0.6%) into the LC, and the fluid intake of 0.5 M sucrose was measured. In both Tg and non-Tg animals, the intake was not significantly different between the PBS- or PhAc-injected groups (***Figure 3D***, n = 4-7 for each group, unpaired two-tailed t-test, t_8_ = 0.061, p = 0.9531 for Tg mice, t_11_ *=* 0.595, p = 0.5636 for non-Tg mice). To further confirm that the activated LC NE neurons does not affect sensitivity to the taste stimulus, we checked the hedonic responses of unconditioned mice to 0.5 M sucrose using the taste reactivity test. Tg mice were given bilateral injections of PBS or 0.6% PhAc into their LC 20 min prior to the intraoral infusion of 0.5 M sucrose. Animal behavior was recorded, and the latency to express the hedonic responses (tongue protrusion and lateral tongue protrusion) was measured. The latency for these responses did not significantly differ between the PBS- and PhAc-injected groups (28.83 ± 8.03 and 33.83 ± 8.33 s, respectively, ***Figure 3E***, n = 6 for each group, unpaired two-tailed t-test, t_10_ *=* 0.432, p = 0.6749), excluding the possibility that the reduction in the latency by LC stimulation simply results from increased sensitivity to the taste stimulus.

It is also possible that the activation of LC NE neurons may promote a state of general arousal, resulting in enhancement of unconditioned responses to aversive events irrespective of memory retrieval. To test this possibility, we examined whether the activation of LC NE neurons alters the rejection responses of mice to an unconditioned bitter tastant, quinine, using the reactivity test. Tg mice received bilateral injection of PBS or 0.6% PhAc into the LC 20 min prior to the intraoral infusion of the 200 μM quinine solution, which induced substantial aversive responses. The latency for rejection responses was not significantly different between the PBS- and PhAc-injected groups (57.75 ± 13.26 s and 54.25 ± 11.34 s, respectively, ***Figure 3F***, n = 8 for each group, unpaired two-tailed t-test, t_14_ = 0.201, p = 0.8439). These data indicate that the activation of LC NE neurons does not enhance unconditioned response to aversive stimulus and exclude the possibility that the reduced latency by LC stimulation simply results from increased arousal level and potentiated general emotional responses to an innate gustatory stimulus.

In addition, locomotor activity of the treated mice was monitored in the open field apparatus. Mice were habituated to the environment for 60 min and injected bilaterally with PBS and then with PhAc solution (0.6%) into the LC. Locomotor activity (60 min) was not significantly different among the Tg and non-Tg groups after PBS or PhAc injection (***Figure 3G***, n = 7 for each group, two-way ANOVA, group effect: F_(1, 12)_ = 0.512, p = 0.4878, time-block effect: F_(17, 204)_ = 60.47, p < 0.0001, and interaction: F_(17,204)_ = 0.480, p = 0.9599). Locomotion of Tg mice after PhAc injection was apparently normal, suggesting that the reduced latency of taste reactivity by LC stimulation cannot be explained by changes in general motor behavior by PhAc injection.

Based on these behavioral evidences, we conclude that the activation of LC NE neurons in response to the exogenous ligand promotes the retrieval process of aversive memory for events acquired through conditioning with a specific taste stimulus.

### Neural pathway mediating enhanced memory retrieval through LC activation

The formation of conditioned taste aversion requires the function of the BLA in the amygdala (***Bermúdez-Rattoni, 2004; Yamamoto et al., 1994***), which receives innervation from LC NE neurons (***Chandler et al., 2014; Robertson et al., 2013***). To determine whether the LC-amygdalar noradrenergic pathway is involved in the enhancement of memory retrieval, we performed pharmacological blockade experiments for taste reactivity using prazosin (PRAZ) and propranolol (PROP) as α_1_- and β-adrenergic receptor antagonists, respectively. First, to check doses of adrenergic receptor antagonists infused into the BLA that do not affect taste reactivity in wild-type mice, we tested the effect of bilateral intra-BLA treatment with PRAZ (0.1 or 0.5 μg/site) or PROP (0.4 or 2.0 μg/site) on the wild-type reactivity. The CS fluid was then intraorally infused, and the latency for reaction response initiation was measured. The latency to reject the CS differed significantly among the groups (***Figure 4A*,** n = 7-10, one-way ANOVA, F_(4, 36)_ = 4.830, p = 0.0032), and the values for the groups treated with higher doses of PRAZ (340.75 ± 38.63 s) and PROP (334.57 ± 34.66 s) were significantly lengthened compared to the PBS-treated controls (156.1 ± 37.42 s, p = 0.0138 for PRAZ, p = 0.0252 for PROP). In contrast, the latency in the groups treated with lower doses of PRAZ (190.75 ± 39.12 s) or PROP (187.75 ± 47.88 s) did not show any significant differences compared to PBS treatment (p = 0.9681 for PRAZ, p = 0.9770 for PROP). Thus, intra-BLA infusion of lower doses of PRAZ and PROP does not influence the latency of taste reactivity in wild-type animals, whereas the data obtained from the use of higher doses of the antagonists show the necessity of the amygdala for the retrieval process of conditioned taste aversion.

**Figure 4.**
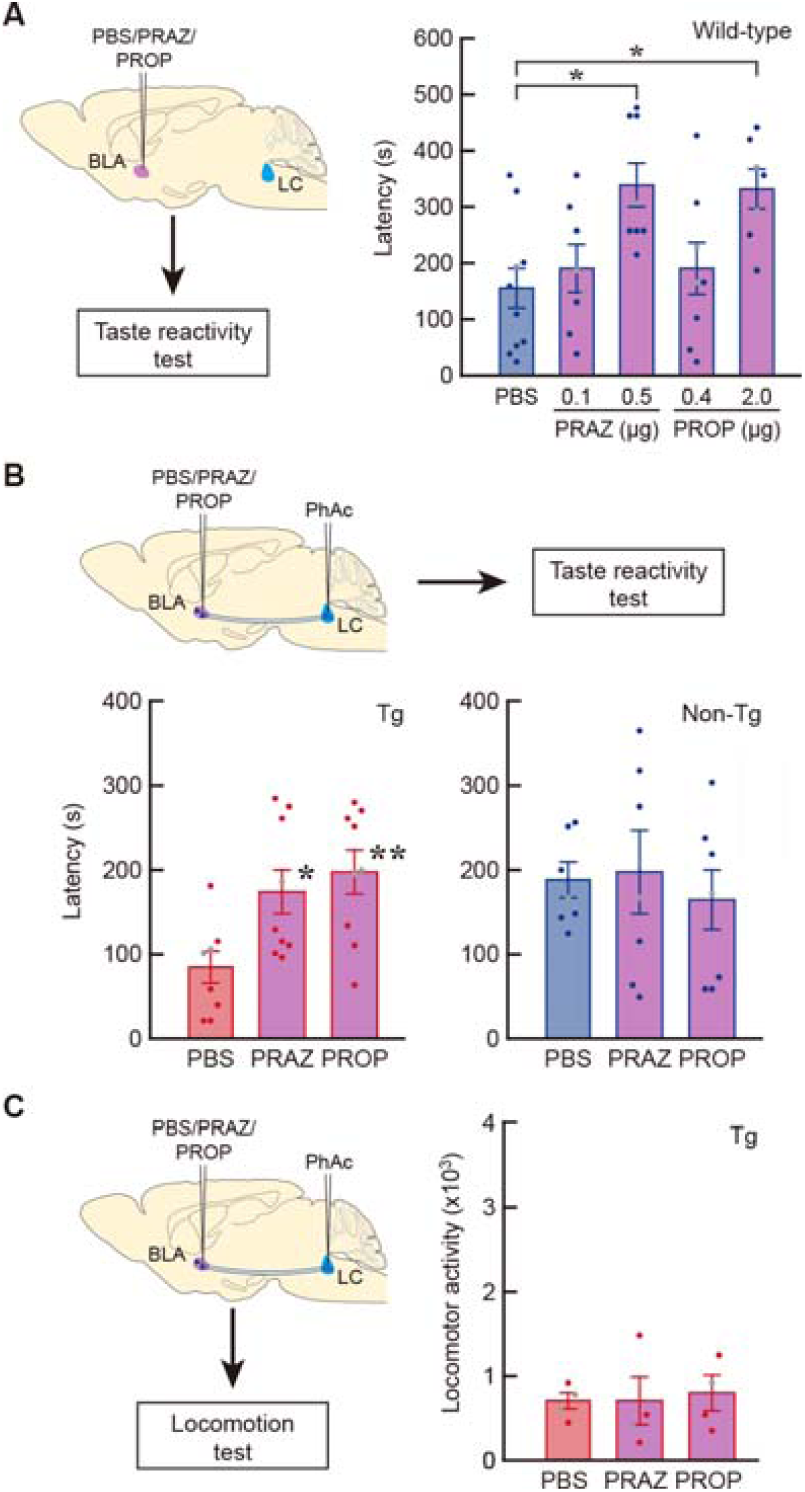
Pharmacological blockade of enhanced memory retrieval by LC activation. (**A**) Taste reactivity test in wild-type mice that received intra-BLA infusion of PRAZ (0.1 or 0.5 μg/site) and PROP (0.4 or 2.0 μg/site). n = 7-10 for each mouse group. *p < 0.05 vs PBS infusion (Tukey HSD test). (**B**) Taste reactivity test showing the blockade of shortened response latency in LC-activated Tg mice by the intra-BLA infusion of PRAZ and POROP at the lower doses (0.1 and 0.4 μg/site, respectively). n = 8 or 9 for each group in Tg mice. n = 7 for each non-Tg group. *p < 0.05, **p < 0.01 vs PBS in Tg mice (Tukey HSD test). (**C**) Locomotion test of the Tg mice that received the microinjection of PhAc (0.6%) into the LC and infusion of PRAZ and POROP (0.1 and 0.4 μg/site, respectively) into the BLA. After the habituation to the open field, the mice received the drug treatments, and then the total number of beam breaks (locomotor activity) during a 60-min period was monitored. n = 4 for each group in Tg mice. Data are presented as mean ± SEM. Individual data points are overlaid.

Next, we tested whether enhanced memory retrieval by LC stimulation can be blocked by the intra-BLA treatment with lower doses of adrenergic receptor antagonists that do not affect taste reactivity in wild-type animals. The Tg and non-Tg mice were given the bilateral intra-BLA treatment of PRAZ (0.1 μg/site) or PROP (0.4 μg/site), which was then followed by the injection of PhAc solution (0.6%) into the LC. The latency to reject the CS in the Tg mice significantly differed among the three groups (***Figure 4B***, n = 8-9 for each group, one-way ANOVA, F_(2, 23)_ = 5.860, p = 0.0088), and the values for the PRAZ- and PROP-treated groups (176.00 ± 26.81 s and 199.11 ± 25.96 s, respectively) were significantly elevated compared to the PBS-treated controls (83.75 ± 19.63 s, p=0.0402 for PRAZ, p=0.0092 for PROP). By contrast, the latency in the non-Tg mice for the treatment of PRAZ (196.14 ± 47.75 s) or PROP (162.57 ± 37.33 s) displayed no significant differences compared to PBS treatment (186.43 ± 20.10 s) (***Figure 4B***: n = 7 for each group, F_(2, 18)_ = 0.220, p = 0.8050). Therefore, the enhanced retrieval of conditioned taste memory by LC noradrenergic activation is blocked by either α_1_- or β-adrenergic antagonist treated into the BLA, suggesting that the retrieval process is mediated, at least in part, through α_1_- and β-adrenergic receptor signalling via the LC-BLA pathway.

In addition, the locomotor activity of the treated mice was monitored in the open field apparatus. The Tg mice were habituated to the environment for 60 min and injected bilaterally with PBS, PRAZ (0.1 μg/site) or PROP (0.4 μg/site) into the BLA, and then PhAc solution (0.6%) into the LC. Locomotor activity (60 min) was not significantly different among the treatment groups (***Figure 4C***, n = 4 for each group, one-way ANOVA, F_(2, 9)_ = 0.0738, p = 0.9294), suggesting that the prolonged latency of taste reactivity by the treatment of adrenergic receptor antagonists into the BLA compared to the PBS injection cannot be attributed to changes in general motor behavior by drug treatment into the BLA.

### Impact of pharmacological inhibition of LC neurons on memory retrieval

To ascertain the enhancing effect of LC noradrenergic activation on memory retrieval, we tested whether the inhibition of LC NE activity would suppress the taste memory retrieval process. Wild-type mice were given an infusion of clonidine (CLO), which is an α_2_ adrenergic receptor agonist that inhibits LC activity (***Aghajanian and VanderMalen, 1982; Washburn and Moises, 1989*)**, into the bilateral LC and subjected to the reactivity test. Latency to reject the CS differed significantly according to the treatments (***Figure 5A***, n = 7-8 for each group, one-way ANOVA, F_(2, 20)_ = 17.19, p < 0.0001), the value for the lower (10 ng/site) or higher (25 ng/site) doses of CLO (380.86 ± 20.13 s and 292.50 ± 29.23 s, respectively) exhibited a significant lengthening compared to PBS injection (165.25 ± 26.14 s, p < 0.0001 for the lower dose, p = 0.0055 for the higher dose). Another cohort of mice that received intra-LC injection of CLO were subjected to the locomotion test, and locomotor activity after the drug treatment was apparently normal, suggesting that CLO treatment does not affect general motor behavior at the habituated condition in the open field (***Figure 5 - figure supplement 1***). The inhibition of LC NE activity actually resulted in the suppression of the retrieval of taste memory, which is consistent with the facilitative role of these neurons in the memory retrieval process.

**Figure 5.**
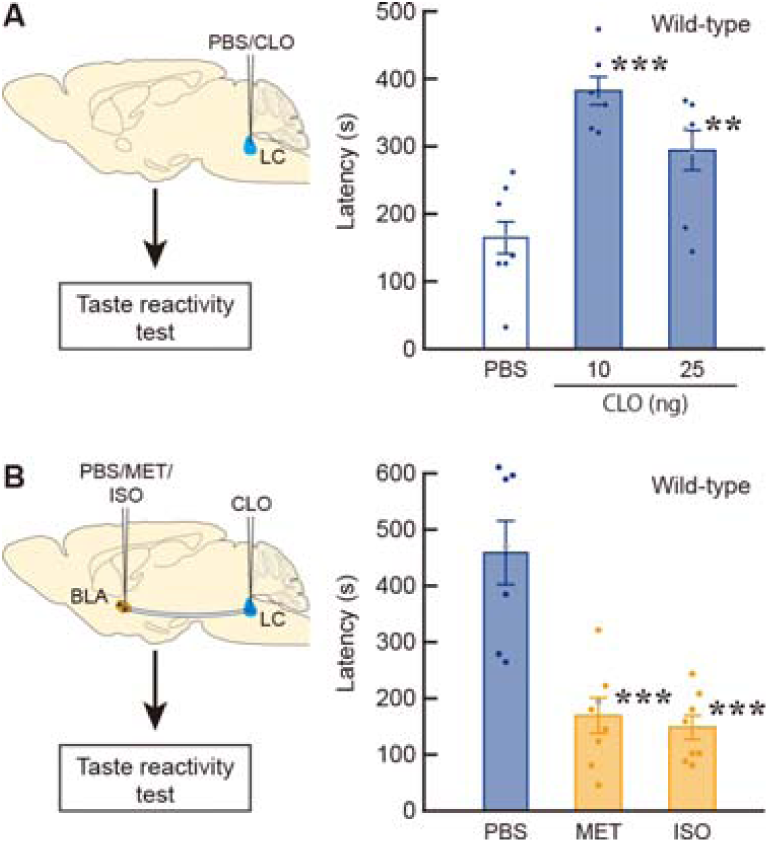
Pharmacological inhibition of LC neurons impairs memory retrieval. (**A**) Taste reactivity test showing the lengthened latency of rejection response in wild-type mice by injection of an α_2_-adrenergic receptor agonist CLO (10 and 25 ng/site) into the LC. n = 7-8 for each mouse group. **p < 0.01, ***p < 0.001 vs the PBS-treated mice (Tukey HSD test). (**B**) Restoration of delayed rejection response in the CLO-injected mice (10 ng/site) by intra-BLA infusion of adrenergic receptor agonists MET (0.5 μg/site) and ISO (1.25 g/site). n = 8 for the MET and ISO-treated groups, n = 7 for PBS-treated group. ***p < 0.001 vs the PBS-treated mice (Tukey HSD test). Data are presented as mean ± SEM. Individual data points are overlaid.

To further validate adrenergic receptor function in the BLA during memory retrieval, we investigated whether the activation of adrenergic receptor subtypes in the amygdala can restore suppressed memory retrieval by inhibition of LC activity. Wild-type mice received bilateral intra-LC infusion of CLO (10 ng/site), and then bilateral intra-BLA treatment of α_1_-adrenergic receptor agonist methoxamine (MET) or β-adrenergic receptor agonist isoproterenol (ISO) prior to the taste reactivity test. The latency to reject the CS in the CLO-injected mice differed significantly according to treatment into the BLA (***Figure 5B***, n = 7-8 for each group, one-way ANOVA, F_(2, 20)_ = 20.83, p < 0.0001), and the values in MET and ISO treatment (169.00 ± 31.06 s and 150.00 ± 21.45 s, respectively) showed a significant shortening compared to PBS treatment (462.00 ± 56.65 s, p < 0.0001 for the MET and ISO treated groups). The data indicate that α_1_- or β-adrenergic receptor activation in the amygdala indeed recovered the impaired memory retrieval by LC inhibition, supporting the idea that the retrieval process of conditioned aversive memory is mediated at least partly through α_1_/β-adrenergic receptor signalling via the LC-BLA pathway.

## Discussion

We successfully developed INTENS as a novel chemogenetic approach to stimulate the activity of neuronal types of interest. The IR84a/IR8a complex forms an odorant-gated ionotropic cation channel (***Abuin et al., 2011; Benton et al., 2009; Grosjean et al., 2011; Rytz et al., 2013***). The present results confirmed that expression of this complex is sufficient to confer PhAl/PhAc-induced excitatory responses in mammalian cells.

Microinjection of PhAc (0.4 or 0.6%) into the LC results in an increased NE release in the Tg brain and that, in particular, a higher dose of PhAc injection was followed by a sustained elevation of NE release in the ACC and the BLA. The IR84a/IR8a-dependent activation generates principally monovalent cation currents (Na^+^ and K^+^), but it may also lead to a small amount of Ca^2+^ entry (***Abuin et al., 2011; Ng et al., 2019***). The intracellular signalling cascades mediated by Ca^2+^ influx may contribute to the sustained elevation of NE release after the high-dose PhAc injection. For instance, Ca^2+^/calmodulin-dependent protein kinase II and protein kinase C mediate trafficking of glutamate receptors and long-term plasticity dependent on protein synthesis (***Herring and Nicoll, 2016; MacDonald et al., 2001***). Activation of protein kinase C also promotes TH gene expression and NE biosynthesis in catecholamine-producing cells (***Goc et al., 1992; Vyas et al., 1990***).

The system of designer receptors exclusively activated by designer drugs has provided a representative chemogenetic strategy that stimulates or inhibits the activity of specific neuronal types (***Roth, 2016; Urban and Roth, 2015***). In this system, the engineered metabotropic receptors are expressed in the target neuronal type and clozapine N-oxide (CNO), a ligand for the engineered receptors, is administered.

However, CNO-induced cellular responsiveness requires G proteins and relevant second messenger cascades that exist endogenously in each neuronal type. In addition, CNO has multiple dose-dependent effects on wild-type animals through *in vivo* conversion to clozapine and N-desmethylclozapine, which act on several types of G protein-coupled receptors (***MacLaren et al., 2016***), and the converted clozapine appears to bind the engineered receptors (***Gomez et al., 2017***). By contrast, our INTENS technology depends on the expression of foreign ligand-gated excitatory IRs, which are directly activated by PhAl/PhAc, in specific neurons, and, basically, does not require other signalling systems. In our technology, a higher dose of ligands (0.6% PhAc) can induce a long-term sustained activation of cellular responsiveness. Furthermore, systemic administration of a ligand precursor enables the stimulation of the neurons in the brain. This technology thus provides a powerful strategy that stimulates the target neuronal activity using the invertebrate-derived ionotropic receptors to study behavioral and physiological roles of neuronal types of interest. Although we used just one type of IR complex here, this family of olfactory receptors represents a large, and functionally divergent, family of ion channel receptors (***Croset et al., 2010***), other members of this group may be useful for different applications of INTENS.

A chemogenetic strategy with chimeric ion channels has been recently reported, in which modified α7 nicotinic acetylcholine receptor ligand-binding domains are fused to an ion pore domain of serotonin receptor 3 (α α (***Magnus et al., 2011, 2019***). The ligand varenicline shows higher affinity to these chimeric receptors as compared to endogenous acetylcholine. In the inhibitory α7-GlyR system, a single systemic injection of the ligand appears to cause sustained behavioral changes for 3 to 4 h, whereas the *in vivo* effects of the stimulatory α7-5HT3 system have not yet been reported. In addition, the inhibitory strategy of neuronal activity has been reported using an ivermectin-sensitive chloride channel (***Lerchner et al., 2007; Slimko et al., 2002***).

In our slice electrophysiology, about 30% of neurons caused depolarization block in response to 0.1% PhAc. Depolarization block is a generally observed phenomenon in excitatory ionic channels. However, this event was not observed in our *in vivo* electrophysiology. In addition, the microdialysis experiment showed that PhAc treatment induced a sustained elevation of NE release in a dose-dependent manner.

These data indicated that *in vivo* treatment of PhAc caused the increased net activity of LC NE neurons in the Tg mice. In our preliminary experiments of the *in vivo* electrophysiology, pneumatic injection of high concentration of glutamate (> 1 mM) frequently induces depolarization block. This suggests that the conditions of PhAc treatment in the present study may have mild excitatory effects, which do not lead to depolarization block, on LC NE neurons expressing IR84a/IR8a receptors.

The present findings provide clear evidence for the key role of LC NE neurons in the retrieval process of conditioned taste aversion, which requires BLA function. A previous study with dopamine β-hydroxylase-deficient mice reports that NE transmission is involved in the retrieval of intermediate-term contextual fear and spatial memory requiring the hippocampus (***Murchison et al., 2004***), whereas pharmacological blockade of β-adrenergic receptors in the BLA does not influence the recall of conditioned odour aversion that depends on the association between an odour stimulus and a visceral malaise-inducing stimulus (***Miranda et al., 2007***). Conditioned odour aversion also requires amygdalar function (***Ferry and Di Scala, 1997***), and the acquisition of this aversion is disrupted by catecholamine depletion of amygdala (***Fernandez-Ruiz et al., 1993***) or blockade of β-adrenergic receptor in the BLA (***Miranda et al., 2007***), suggesting a distinct mechanism on the maintenance or retrieval of conditioned aversive memory between sensory modalities. In addition, electrical LC stimulation facilitates memory retrieval of the maze performance (***Devauges and Sara, 1991; Sara and Devauges, 1988***), although the target brain regions affected by LC activity have not been identified. A functional magnetic resonance imaging study also indicates the LC response at the retrieval of events learned in an emotional context in humans (***Sterpenich et al., 2006***). These data suggest the engagement of LC NE neurons in the retrieval process of different types of memory.

LC NE neurons send their axon terminals to nuclei in the amygdala, including the BLA and central nucleus (CeN) (***Chandler et al., 2014; Robertson et al., 2013***).

Conditioned taste aversion depends on the amygdala function, particularly the BLA (***Bermúdez-Rattoni, 2004; Yamamoto et al., 1994***). In the present study, the coordinates used for the drug injection into the amygdala were localized within the BLA. Our results show that the facilitative role of LC NE neurons in the retrieval process of taste aversion was mediated at least partly through α_1_- and β-adrenergic receptors via the LC-BLA pathway. Although some studies indicate the role of the CeN in taste aversion (***Baha et al., 2003; Lamprecht et al., 1997***), the requirement of the CeN for the learning is controversial (***Yamamoto et al., 1995***). It remains unclear whether LC projection to the CeN is involved in the retrieval process of the taste aversion task.

A recent study reported that the BLA is required for the retrieval process of conditioned taste aversion (***Inui et al., 2019***). The LC-BLA pathway may directly influence the activity of BLA neurons at the phase of memory retrieval of taste aversion. The insular cortex is also required for the formation of taste aversion (***Bermúdez-Rattoni, 2004; Yamamoto et al., 1994***) and especially for long-term memory formation dependent on *de novo* protein synthesis and mitogen-activated protein kinase cascades (***Berman et al., 1998; Rosenbluk.et al., 1993***). These data suggest the engagement of the insular cortex to memory consolidation through interaction with the amygdala. This evidence suggests that LC noradrenergic neurons influence the interaction between the cortical and amygdala regions to facilitate the recall of stored memory.

NE terminals are localized in both pyramidal neurons and GABAergic interneurons in the BLA (***Farb et al., 2010; Li et al., 2001, 2002***). About half of the number of NE axons form synapses on dendritic shafts and spines of pyramidal neurons, and a small number of the axons make synapses on cell bodies and dendrites of presumptive interneurons (***Zhang et al., 2013***). NE potentiates glutamate-mediated excitatory synaptic transmission through presynaptic mechanisms via β-adrenergic receptors in the amygdala (***Ferry et al., 1997; Huang et al., 1996***). β-Adrenergic receptor stimulation appears to enhance spike frequency in pyramidal neurons in the amygdala of juvenile mice (***Fink and LeDoux, 2018***). NE facilitates GABA transmission through presynaptic mechanisms via α1-adrenergic receptors (***Braga et al., 2004***), and also excites some GABA interneurons directly via α_1_-adrenoceptors (***Kaneko et al., 2008***). These potential synaptic mechanisms may underlie the facilitation of the retrieval process of taste aversion memory through the activation of the LC-BLA pathway. *In vivo* electrophysiological studies have reported that LC stimulation or NE iontophoresis inhibits spontaneous firing of the majority of BLA neurons and decreases the responsiveness of these neurons to cortical stimulation in normal conditions (***Buffalari and Grace, 2007; Chen and Sara, 2007***). However, in response to chronic stress exposure, NE increases spontaneous activity of BLA neurons and produces a facilitation of responses evoked by cortical stimulation (***Buffalari and Grace, 2009)***. These data suggest a dynamic shift in the NE response of amygdala neurons dependent on the environmental conditions or emotional context. Therefore, it is necessary to elucidate the detailed mechanisms underlying the synaptic mechanisms through adrenergic receptor subtypes in different types of BLA neurons to facilitate the retrieval process in conditioned animals. In addition, recent studies report that dopamine is released from LC terminals in the hippocampus and that LC-derived dopamine modulates memory formation (***Kempadoo et al., 2016; Takeuchi et al., 2016***).

Similarly, dopamine may be released from LC terminals in the amygdala and partly implicated in the retrieval process of amygdala-dependent memory. This issue needs to be addressed in the future.

In conclusion, the present study provides evidence that the LC-BLA pathway, through α1- and β-adrenergic receptor signalling, is essential and sufficient for enhancing the retrieval process of conditioned taste aversion memory. We focused on the study of LC NE function in taste aversion circuits, and we need to investigate whether the finding on the role of these neurons in memory recall is applicable for other types of memory paradigms. Dysfunction of the central NE system is implicated in the pathological states of neuropsychiatric diseases, such as Korsakoff’s syndrome with anterograde and retrograde amnesia, post-traumatic amnesia, and post-traumatic stress disorder (***Chamberlain and Robbins, 2014; Hendrickson and Raskind, 2016***).

Adrenergic receptor subtypes are potential therapeutic targets for post-traumatic stress disorder (***Strawn and Geracioti, 2008***). A detailed analysis of the neural substrate that mediates memory retrieval dependent on the LC-amygdalar NE system may provide insight leading to a deeper understanding of the mechanisms underlying the pathogenesis of these neuropsychiatric diseases.

## Contact for resources and reagents sharing

Further information and requests for resources and reagents should be directed to and will be fulfilled by the Lead Contact, Kazuto Kobayashi.

## Materials and methods

### Animals

A gene cassette encoding a chimeric protein composed of GFP-IR84a-2A-IR8a with a human calreticulin signal peptide was exchanged by the GFP cDNA part of the plasmid pTH-GFP that contains a 9-kb rat TH gene promoter, rabbit β-globin second intron, GFP cDNA, and rabbit β -globin and SV40 early gene polyadenylation signals (***Sawamoto et al., 2001; Matsushita et al., 2002***), resulting in the plasmid pTH-GFP-IR84a/IR8a. The transgene construct was linearized by *Sal*I digestion, purified by a gel electrophoresis, and microinjected into fertilized C57BL/6J mouse eggs, which were then implanted into pseudopregnant females. Tg mice were identified by Southern blot hybridization or polymerase chain reaction with genomic DNA prepared from tail clips.

We generated 25 independent transgenic founders carrying the TH-EGFP-IR84a/IR8a transgene. The transgenic offspring derived from each founder were subjected to GFP immunostaining with sections prepared from the transgenic brain. Expression levels and distribution of the transgene in the brain regions including the olfactory bulb, ventral midbrain, and hindbrain, showed variations among the strains. Based on the specificity and level of transgene expression, we selected one transgenic line, the TH-EGFP-IR84a/IR8a-2-1 line. This strain was used for the following experiments in the present study.

Mice were maintained on a 12 h light/dart cycle (lights on at 07:00 h), at an ambient temperature of 22℃. All experimental procedures were conducted during the light period. The mice aged 12-14 weeks old were used for the following experiments except for *in vitro* electrophysiology, in which the animals aged 17-20 postnatal days were used. The mice were housed in groups of three to five, and they were singly housed after the stereotaxic surgeries for the microdialysis and behavioral experiments. Assignment of animals to the experimental conditions was random.

### Histology

Mice were anesthetized with sodium pentobarbital (50 mg/kg, intraperitoneal) and perfused transcardially with PBS, followed by fixation with 4% paraformaldehyde in 0.1 M phosphate buffer (pH 7.4). Sections (30 μm thick) were incubated with a primary antibody for GFP (rabbit, 1:2,000, Invitrogen) and then with a biotinylated secondary antibody (anti-rabbit IgG, 1:500, Jackson ImmunoResearch Laboratories). The immunoreactive signals were visualized by use of a Vectastain Elite ABC kit (Vector Laboratories) with 3,3C-diaminobenzidine tetrahydrochloride/H_2_O_2_ as a chromogen. For double-fluorescence histochemistry, sections were incubated with anti-TH antibody (mouse, 1:400, Millipore) and anti-GFP antibody (rabbit, 1:2,000, ThermoFisher) or anti-NET antibody (mouse, 1:2,000) and anti-IR8a antibody (rabbit, 1 μg/ml). Anti-IR8a antibody was prepared by immunization with purified protein of glutathione-S-transferase fused to a 27-amino acid fragment of IR8a (DKYSPYSSRNNRQAYPVACREFTLRES). The specificity of anti-IR8a antibody was confirmed by double-fluorescence immunohistochemistry for NET/IR8a with LC sections prepared from the Tg and non-Tg mice. The sections were incubated with species-specific secondary antibodies conjugated to Alexa488 (Molecular Probes) and Cy3 (Jackson ImmunoResearch). Fluorescent images were visualized under a confocal laser-scanning microscope (LSM510 or LSM800, Zeiss) equipped with proper filter cube specifications.

For cell counts of double-fluorescence histochemistry for TH and GFP, 4 sections through the LC along with the anteropoterior coordinates (mm) between −5.43 and −5.68 from bregma were prepared from each mouse and used for double immunostaining. The number of immuno-positive cells in the region of interest (200 × 200 μm) was counted by using a computer-assisted imaging program (NIH Image 1.62, National Institutes of Health), and the number of TH^+^/GFP^+^ cells was divided by that of total TH^+^ cells in each section. The average of the percentage obtained from 4 sections was calculated. Four mice were used for cell counts of the immunostaining.

For *in situ* hybridization, Fresh-frozen sections (10 μm thick) were fixed in a solution of 4% paraformaldehyde in 0.1 M phosphate buffer and treated with 0.1 M triethanolamine (pH 8.0) containing 0.25% acetic anhydride. Sections were hybridized with antisense RNA probe for IR84a (nucleotides 82-780) or IR8a (nucleotides 4-529) sequence labelled using *in vitro* transcription with digoxigenin-11-UTP (Roche). The signals were visualized with a nonradioactive detection system using anti-digoxigenin Fab fragment conjugated to alkaline phosphatase (Roche).

To validate the cytotoxicity of PhAc treatment, sections through the LC were prepared from the Tg mice 7 days after unilateral treatment with PBS or PhAc (0.4/0.6%) for the microdialysis analysis, and stained for TH and NET immunohistochemistry. The ratio of the number of cells stained for TH or NET in the treated side relative to the intact side was calculated. In the experiment for the staining with cell death markers, ibotenic acid (IBO, 1 mg/ml, as positive controls to detect cell death signals) or PhAc (0.6%) was injected into the LC (0.2 μl/site) in either hemisphere of the Tg mice. The LC sections were stained with terminal deoxynucleotidyl transferase-mediated dUTP-biotin nick end labeling (TUNEL) or immunostaining for activated caspase-3, together with counter staining with 4’,6-diamidino-2-phenylindole (DAPI).

### Electrophysiology

For slice electrophysiology, mice were anesthetized with 1.5% isoflurane. Coronal brain slices containing the LC were cut (300 μm thick) using a microslicer (PRO7, Dosaka) in ice-cold oxygenated cutting Krebs solution of the following composition (mM): choline chloride, 120, KCl, 2.5, NaHCO_3_, 26, NaH_2_PO_4_, 1.25, D-glucose, 15, ascorbic acid, 1.3, CaCl_2_, 0.5, and MgCl_2_, 7. The slices were then transferred to a holding chamber containing standard Krebs solution of the following composition (mM): NaCl, 124, KCl, 3, NaHCO_3_, 26, NaH_2_PO_4_, 1, CaCl_2_, 2.4, MgCl_2_, 1.2, and D-glucose, 10 (pH 7.4) when bubbled with 95% O_2_-5% CO_2_. Slices were incubated in the holding chamber at room temperature (21°-26°C) for at least 1 hour before recording. Neurons in the LC were visualized with a 60× water immersion objective attached to an upright microscope (BX50WI, Olympus Optics). LC neurons with fluorescence were visualized using the appropriate fluorescence filter (U-MWIG3, Olympus Optics). Patch pipettes were made from standard-walled borosilicate glass capillaries (Harvard Apparatus). For the recording of membrane potentials, a K-gluconate-based internal solution of the following composition (mM) was used: K-gluconate, 120, NaCl, 6, CaCl_2_, 5, MgCl_2_, 2, K-EGTA, 0.2, K-HEPES, 10, Mg-ATP, 2, and Na-GTP, 0.3 (pH adjusted to 7.4 with 1 M KOH). Whole-cell recordings were made from LC neurons with fluorescence using a patch-clamp amplifier (Axopatch 200B, Molecular Devices). Data were stored on digital audiotapes using a DAT recorder (DC to 10 kHz, Sony), and were digitized off-line at 10 kHz (low-pass filtered at 2 kHz with an 8-pole Bessel filter) using pCLAMP9 software (Molecular Devices).

NE cells in the LC were identified by low frequency (<7 Hz) action potentials with large after-hyperpolarization, as described in previous studies (***van den Pol, et al., 2002; Zhang et al., 2010***). The effects of drugs on the membrane potential were assessed after they had reached a steady state (starting point), and the mean firing frequency and membrane potential were calculated during a 30-s test period before the drug application (pre) and after the starting point (post). Although a few neurons had firing frequency greater than 7 Hz (9-10 Hz) before drug application, these neurons were also included in the analyses due to the characteristic shape of action potentials with large after-hyperpolarization.

For cultured cell electrophysiology, HEK293T cells were transduced by a lentiviral vector carrying a fusion gene that encodes EGFP-IR84a and hemagglutinin (HA)-IR8a connected through 2A peptide under the control of a cytomegalovirus gene promoter. The ligand-induced currents were measured at room temperature using the standard whole-cell voltage-clamp technique (***Osanai et al., 2006; Tarradas et al., 2013***). Borosilicate glass pipettes (4-6 M Ω) were filled with an intracellular solution containing (mM): 135 K-methanesulfonate, 5 KCl, 0.5 EGTA, 5 Mg-ATP, 0.4 Na-GTP, and 10 HEPES (pH 7.2, adjusted with KOH), and were used for voltage clamp recordings. Cells were continuously perfused with a modified Hanks’ solution containing (mM): 137 NaCl, 3 KCl, 1.2 MgCl_2_, 1.8 CaCl_2_, 20 glucose, and 10 HEPES (pH 7.4, adjusted with NaOH after dissolving PhAc). Currents were recorded with an EPC-10 amplifier (HEKA), and the data were sampled at 2 kHz. Non-transduced GFP-negative cells were used for the controls. To monitor ligand dose responses, the amplitudes of the ligand-induced currents were measured at different doses of PhAc and normalized in each cell to the amplitude at the average of 0.01% PhAc treatment.

An extracellular single-unit recording was carried out *in vivo* (***Takahashi et al., 2010***). Mice were anesthetized with 1.5% isoflurane and anaesthesia was maintained with 0.5-1.0% isoflurane based on the monitoring of electroencephalogram. The mice were placed in the stereotaxic frame (SR-5M, Narishige) with ear bars and a mouth- and-nose clamp. Body temperature was maintained at 37-38°C with a heating pad. The scalp was opened, and a hole was drilled in the skull above the LC with the coordinates in mm anteroposterior (AP) −1.2 and mediolateral (ML) +0.9 from lambda according to the mouse stereotaxic atlas (***Franklin and Paxinos, 2008***). Two skull screws were placed over the occipital bone and another skull screw was placed on the frontal bone. One of the two screws over the occipital bone was used as a reference for the recording. For pneumatic injection, the firing activity was recorded by a double-barrel glass pipette, in which the tip of the injection capillary (tip diameter: 50μm, A&M systems) m above the tip of the recording electrode (tip diameter: 2-3 μm, impedance: 15-20 MΩ, Harvard Apparatus). The pipette was lowered into the LC, dorsoventral (DV) −2.2 to −3.5 mm from dura, and recording was performed using Spike2 (Cambridge Electronic Design) at a sampling rate of 20 kHz. NE neurons were identified based on a slow tonic firing (< 7 Hz) with long spike duration (> 0.8 ms) as described (***Takahashi et al., 2010***). Solution of 1% PhAc was delivered into the LC through a pneumatic pump (PV-820, World Precision Instruments). In this technique, a small amount of ligand solution (estimated at 4–10 nl) was infused through the pump (20–40 psi, 10–100 ms). The mean firing rate was calculated for each neuron during a 30-s test period immediately before injection (pre) and after injection (post), and used for comparisons. For systemic administration of ligand precursor, MPhAc was dissolved in PBS containing 1.5% Tween 80 and 2.5% ethanol. Solution of MPhAc (20 mg/kg) or vehicle (5 ml/kg) was administered in the lateral tail vein and the *in vivo* recording was performed as described above, except for the use of the recording electrode without the injection capillary. Baseline firing rate (pre) was recorded during a 3-min test period immediately before systemic administration MPhAc solution. Ten mins after the drug administration, the effects were assessed during 3–10-min test periods (post). After the electrophysiological experiments, the recording sites were marked by iontophoretic injection of 2% pontamine sky blue. The mice were deeply anesthetized with sodium pentobarbital and perfused transcardially with PBS, followed by 10% formaline. Brain sections were stained with neutral red for the verification of recording sites. The post-mortem histological analysis verified deposits of recording sites in the LC (***Figure 2 — figure supplement 2A***).

### Microdialysis

Mice were anesthetized with 1.5% isoflurane and underwent unilaterally stereotaxic surgery with a 25-gauge guide cannula (Eicom) for a dialysis probe aimed at ACC or the amygdala as well as a 30-gauge guide cannula (Eicom) for injection into the LC at the ipsilateral side. The coordinates (mm) from bregma or dura were AP +0.7, ML +0.3, and DV −0.4 for the ACC dialysis probe, AP −1.2, ML +3.5, DV −3.5 for the amygdala dialysis probe, and AP −0.75, ML +0.65, and DV −2.5 for the LC microinjection, according to the mouse stereotaxic atlas (***Franklin and Paxinos, 2008***). Two or three days later, the stylet in the cannula was replaced with an active membrane dialysis probe (1.0 mm in length, 0.22 mm in outer diameter, Eicom) that was connected to a l μ syringe filled with artificial cerebrospinal fluid (aCSF) of the following composition (mM): NaCl, 148, KCl, 4.0, MgCl_2_, 0.85, and CaCl_2,_ 1.2. For the intra-LC injection, the stylet in the LC cannula was replaced with a 35-gauge internal cannula (1 mm beyond the tip of the implanted guide cannula, Eicom) connected to a 10 μl Hamilton syringe, aCSF was pumped through the probe at a rate of 1.0 μl/min for 2 h, and then dialysis samples were collected every 30 min using a refrigerated fraction collector (EFC-82, Eicom). Each sample vial contained 10 μl of antioxidant, which consisted of 20 mM phosphate buffer including 25 mM EDTA-2Na and 0.5 mM of ascorbic acid (pH 3.5). Three baseline samples were collected to measure a tonic level of NE. This was followed by PhAc (0.4/0.6%) or PBS injection into the LC for 5 min to a total volume of 0.2 μl. Four samples were collected thereafter to assess the time course for change in NE level after the injection. The amount of NE in each fraction was determined by a high-performance liquid chromatography system (CA-50DS, 2.1 mm × 150 mm, Eicom, with the mobile phase containing 5% methanol in 100 mM sodium phosphate buffer, pH 6.0) equipped with an electrochemical detector (ECD-300, Eicom). Results are expressed as percentage of baseline concentration (analyte concentration × 100/mean of the three baseline samples). Systemic administration of MPhAc (10 mg/kg) or vehicle (5 ml/kg) was administered in the lateral tail vein and microdialysis samples were collected from the ACC as described above. After the microdialysis experiments, the mice were deeply anesthetized with sodium pentobarbital and perfused transcardially with PBS, followed by 10% formaline. Brain sections were stained with neutral red for the verification of placement sites. The post-mortem analysis confirmed the placement sites of the injection needle in the LC and dialysis probes in the ACC and the amygdala (***Figure 2 — figure supplement 2B-D***).

### Behavioral analysis

Taste reactivity test was carried out as described (***Inui et al., 2013; Yasoshima and Shimura, 2017***) with some modifications. Mice were anesthetized with 1.5% isoflurane and, surgically, bilaterally implanted with 26-gauge guide cannulae (Plastics One) into the LC using the coordinates (mm) AP −0.9, ML +0.6 and DV −2.3 from lambda or dura according to the brain atlas (***Franklin and Paxinos, 2008***). An intraoral catheter for infusion of the taste stimulus (0.5 M sucrose dissolved in tap water) was surgically placed on the left side of the oral cavity, and the inlet of the catheter was fixed with acrylic cement on the guide cannula assembly. After a 1-week recovery period, the mice were placed on a 20-h water deprivation schedule. Fluid intake training with tap water was conducted over 4 days. Consumption was measured by weighing bottles before and after the 20-min access. The mice were allowed to access water *ad libitum* for 3 h after the training. The taste aversion paradigm contained three different phases (***Figure 3A***). In the conditioning phase (days 1 and 2), the mice were presented to 0.5 M sucrose with the spout as the CS for 20 min, and 10 min later they received an intraperitoneal injection with 0.15 M LiCl (2% of body weight) as the US. In the habituation phase (days 3-5), the mice were habituated to intraoral infusion with tap water in the test chamber. The chamber consisted of an acrylic cylinder (dimensions in mm: diameter, 140, height, 250) and an acrylic box (dimensions in mm: length and width, 325, height, 310) equipped with an inside mirror, which enabled monitoring of the animal’s behavior from the bottom, and the infusion was carried out at the constant flow rate of 50 μl/min for 15 min by syringe pump (Eicom). On the day of the test phase (day 6), a solution (0.2 μl/site) containing PhAc (0.4/0.6%) or PBS was injected into the bilateral LC 20 min before CS presentation. The solution was delivered through a 33-gauge internal cannula (1 mm beyond the tip of the implanted guide cannula, Plastic One) at a flow rate of 0.05 μl/min. The mice were placed in the test chamber and infused intraorally with 0.5 M sucrose. Behavior was recorded using a digital video camera, and the latency for the initiation of rejection responses (including gaping, chin rubbing, forelimb flailing, paw wiping, and CS dropping) was measured.

For the taste sensitivity test, mice received a bilateral implantation of guide cannulae into the LC. The mice were placed on a 20-h water deprivation schedule and conducted the intake training over 4 days. They received microinjection of PBS or 0.6% PhAc into the bilateral LC (0.2 μl/site) 20 min before the fluid presentation, 0.5 M sucrose was presented to the mice for 20 min, and the sucrose consumption was measured.

To monitor hedonic responses, mice were surgically bilaterally implanted with 26-gauge guide cannulas into the LC, and then with an intraoral catheter for infusion of the taste stimulus on the left oral cavity as described above. The mice were habituated to the intraoral infusion with tap water in the test chamber. A solution (0.2 μl/site) containing PBS or 0.6% PhAc was injected into the bilateral LC, 20 min later, the mice were placed in the test chamber and infused intraorally with 0.5 M sucrose. Behavior was recorded using a digital video camera, and the latency for the initiation of hedonic responses (including tongue protruding and lateral tongue protruding) was measured.

To test aversive responses to quinine, mice received a bilateral implantation of guide cannulae into the LC and a unilateral placement of an intraoral catheter into the oral cavity. The mice were habituated to the intraoral infusion with tap water in the test chamber. A solution (0.2 μl/site) containing PBS or 0.6% PhAc was injected into the bilateral LC, 20 min later, the mice were placed in the test chamber and infused intraorally with 0.2 mM quinine. Behavior was recorded, and the latency for the initiation of rejection responses was measured.

Prior to the pharmacological blocking experiment with adrenergic receptor antagonists, doses of adrenergic receptor antagonists infused into the amygdala were checked that do not affect the taste reactivity in wild-type mice. The mice were implanted bilaterally with guide cannulae into the amygdala using the coordinates (AP −1.2, ML +3.5, DV −2.5 from bregma or dura) according to the mouse brain atlas (***Franklin and Paxinos, 2008***) and one intraoral catheter on the oral cavity. A solution (0.2 μl/site) containing PRAZ (0.1 or 0.5 μg/site) or PROP (0.4 or 2.0 μg/site) was injected bilaterally through an internal cannula into the amygdala, and then the taste reactivity test was performed as described above. For the blocking experiment, mice were further implanted surgically with two guide cannulae into the bilateral LC using the same coordinates as above. A solution (0.2 μl/site) containing PRAZ (0.1 μg/site) or PROP (0.4 μg/site) was injected bilaterally through an internal cannula into the amygdala 5 min before bilateral injection of 0.6% PhAc into the LC (0.2 μl/site) for the taste reactivity test.

For the pharmacological inhibition experiment of LC neuronal activity, mice were implanted surgically with two guide cannulae into the bilateral LC and one intraoral catheter into the oral cavity as above. Solution (0.2 μl/site) containing CLO (10 and 25 ng/site) was injected bilaterally through an internal cannula into the LC and tested for taste reactivity 20 min later. For the pharmacological recovery, the mice were further implanted bilaterally with guide cannulae into the amygdala using the same coordinates as above. A solution (0.2 μl/site) containing MET (0.5 μg/site) or ISO (1.25 μg/site) was injected bilaterally through an internal cannula into the amygdala 5 min after the bilateral intra-LC injection of CLO (10 ng/site) for the taste reactivity test.

Locomotor activity was measured with a movement analyser equipped with photo-beam sensors (SV-10, Toyo Sangyou). The number of beam breaks was counted for every 10-min session. The total number of beam breaks during a 60-min test period was calculated to evaluate the spontaneous locomotor activity during the pretreatment (sessions −6 to −1) and the locomotor activity after PBS treatment (sessions 1-6) and 0.6% PhAc treatment (sessions 7-12).

Animal behaviors were scored by the experimenters, who were blind to the treatment of the animals. After all behavioral testing, mice were anesthetized with sodium pentobarbital and perfused transcardially with PBS, followed by 10% formaline. Brain sections were stained with cresyl violet for the verification of placement sites.

Histological examination confirmed the placement sites of internal cannulae within the target brain regions (***Figure 3 — figure supplement 2*** for the taste reactivity, sensitivity, and locomotor activity tests, ***Figure 4 — figure supplement 1*** for the pharmacological blocking experiments, and ***Figure 5 — figure supplement 1*** for LC inhibition experiments).

### Quantification and statistical analyses

Although no statistical methods were used to predetermine the sample size for each measure, we employed similar sample sizes to those that were reported in previous publications from our labs, which are generally accepted in the field. In each testing, we conducted an experiment in two to five separate cohorts, each of which was designed to include all experimental groups. Data obtained across cohorts were pooled, and outliers were defined as data points located outside the range of mean ± 2SD. All statistical analyses were two-tailed and conducted using SPSS ver. 25 (IBM). The reliably of the results was assessed against a type I error (α) of 0.05. Prior to unpaired t-test, we assessed equality of variances for two groups with Levene’s test, and if this was not the case, we reported the statistics (F and p), and then corrected the degree of freedom according to Welch’s method. For significant main effects identified in one- and two-way analyses of variant (ANOVAs), post hoc comparisons were performed with Tukey HSD test. For significant interactions revealed in two-way ANOVAs, t-tests with Holm-Bonferroni sequential correction were used post hoc. One, two, and three stars (*) in figures represent p-value < 0.05, 0.01, and 0.001, respectively.

## Acknowledgments

This work was supported by grants-in-aid for Scientific Research on Innovative Areas *Adaptive Circuit Shift* (26112002) from the Ministry of Education, Science, Sports, and Culture of Japan, and Core Research for Evolutional Science and Technology (JP16gm0310008) of Japan Science and Technology Agency (K.K.), and by an ERC Consolidator Grant (615094), an HFSP Young Investigator Award (RGY0073/2011), the SNSF Nano-Tera Envirobot project (20NA21_143082) (R.B.). We are grateful to M. Kikuchi, N. Sato, H. Hashimoto, and M. Sugawara for technical support during animal experiments and to T. Kobayashi for helpful illustrations.

## Additional Information

### Author contributions

**Ryoji Fukabori,** Conceptualization, Data curation, Formal analysis, Investigation, Visualization, Writing-original draft, Writing-review and editing; **Yoshio Iguchi,** Conceptualization; Data curation, Formal analysis, Investigation, Visualization, Writing-original draft, Writing-review and editing; **Shigeki Kato**, Data curation, Investigation, Resources, Visualization, Writing-original draft; **Kazumi Takahashi**, Data curation, Formal analysis, Investigation, Visualization, Writing-original draft; **Satoshi Eifuku**, Data curation, Formal analysis, Investigation, Visualization, Writing-original draft; **Shingo Tsuji**, Data curation, Formal analysis, Investigation, Visualization, Writing-original draft; **Akihiro Hazama**, Data curation, Formal analysis, Investigation, Visualization, Writing-original draft; **Motokazu Uchigashima**, Data curation, Investigation, Resources, Visualization, Writing-original draft; **Masahiko Watanabe**, Data curation, Investigation, Resources, Visualization, Writing-original draft; **Hiroshi Mizuma**, Data curation, Formal analysis, Investigation, Visualization, Writing-original draft; **Yilong Cui**, Data curation, Formal analysis, Investigation, Visualization, Writing-original draft; **Hirotaka Onoe**, Data curation, Formal analysis, Investigation, Visualization, Writing-original draft; **Keigo Hikishima**, Data curation, Formal analysis, Investigation, Visualization, Writing-original draft; **Yasunobu Yasoshima**, Data curation, Formal analysis, Investigation, Visualization, Writing-original draft; **Makoto Osanai**, Data curation, Formal analysis, Investigation, Visualization, Writing-original draft; **Ryo Inagaki,** Data curation, Formal analysis, Investigation, Visualization, Writing-original draft; **Kohji Fukunaga**, Data curation, Formal analysis, Investigation, Visualization, Writing-original draft; **Takuma Nishijo**, Data curation, Formal analysis, Investigation, Visualization, Writing-original draft; **Toshihiko Momiyama**, Data curation, Formal analysis, Investigation, Visualization, Writing-original draft; **Richard Benton**, Conceptualization, Funding acquisition, Resources, Supervision, Writing-original draft; **Kazuto Kobayashi**, Conceptualization, Funding acquisition, Resources, Supervision, Validation, Visualization, Writing-original draft, Writing-review and editing.

### Author ORCIDs

Ryoji Fukabori https://orcid.org/0000-0001-7534-8237

Yoshio Iguchi https://orcid.org/0000-0001-8240-3345

Shigeki Kato https://orcid.org/0000-0002-8792-7591

Kazumi Takahashi https://orcid.org/0000-0003-4015-8016

Satoshi Eifuku https://orcid.org/0000-0003-0032-1720

Akihiro Hazama https://orcid.org/0000-0002-1199-8246

Motokazu Uchigashima https://orcid.org/0000-0002-0878-2233

Masahiko Watanabe https://orcid.org/0000-0001-5037-7138

Hiroshi Mizuma https://orcid.org/0000-0001-8970-9486

Yilong Cui https://orcid.org/0000-0002-8302-1899

Keigo Hikishima https://orcid.org/0000-0003-1133-1297

Makoto Osanai https://orcid.org/0000-0002-3572-8195

Ryo Inagaki https://orcid.org/0000-0002-0996-5269

Kohji Fukunaga https://orcid.org/0000-0001-8526-2824

Takuma Nishijo https://orcid.org/0000-0003-0533-0132

Toshihiko Momiyama https://orcid.org/0000-0002-3588-9167

Richard Benton https://orcid.org/0000-0003-4305-8301

Kazuto Kobayashi https://orcid.org/0000-0002-7617-2939

### Ethics

Animal care and handling procedures were conducted in accordance with the guidelines established by the Experimental Animal Centre of Fukushima Medical University. All procedures were approved by the Fukushima Medical University Institutional Animal Care and Use Committee.

## Additional files

### Supplementary files

Transparent reporting form

## Data availability

The datasets generated and analysed in this study are deposited on Mendeley data. DOI: 10.17632/fbgbrbjz6z.3

## Figure Supplements

**Figure 1 - figure supplement 1.**
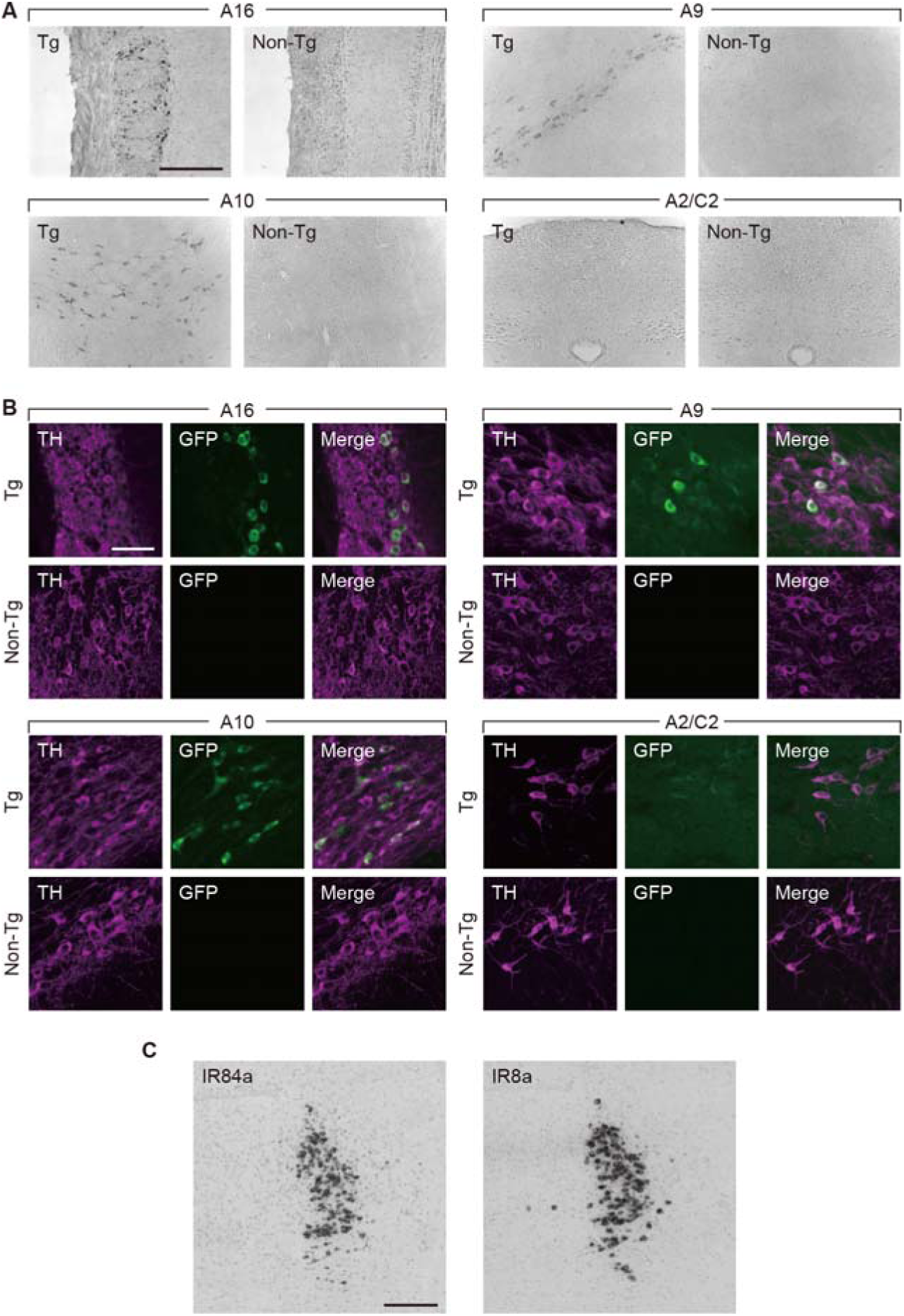
Histological analysis of transgene expression. (**A**) Expression of transgene in catecholamine-containing cell groups in TH-EGFP-IR84a/IR8a mice. Sections through the olfactory bulb (A16), ventral midbrain (A10 and A9), and brain stem (A2/C2) of Tg and non-Tg mice were prepared and stained by GFP immunohistochemistry. Transgene expression was observed in many cells in the A16, A10, and A9 cell groups (dopamine neurons) and in a few cells in the A2/C2 cell groups (NE/epinephrine neurons) of the Tg mice. (**B**) Double immunostaining for TH/GFP of brain sections through the A16, A10, A9, and A2/C2 cell groups. Confocal microscopic images of the sections are shown. (**C**) In situ hybridization showing expression of IR84a or IR8a mRNAs in Tg mice. Microscopic images of LC sections obtained from *in situ* hybridization with the antisense probe for IR84a or IR8a. Scale bars: 200 μm (A); 50 μm (B); 400 μm (C).

**Figure 1 - figure supplement 2.**
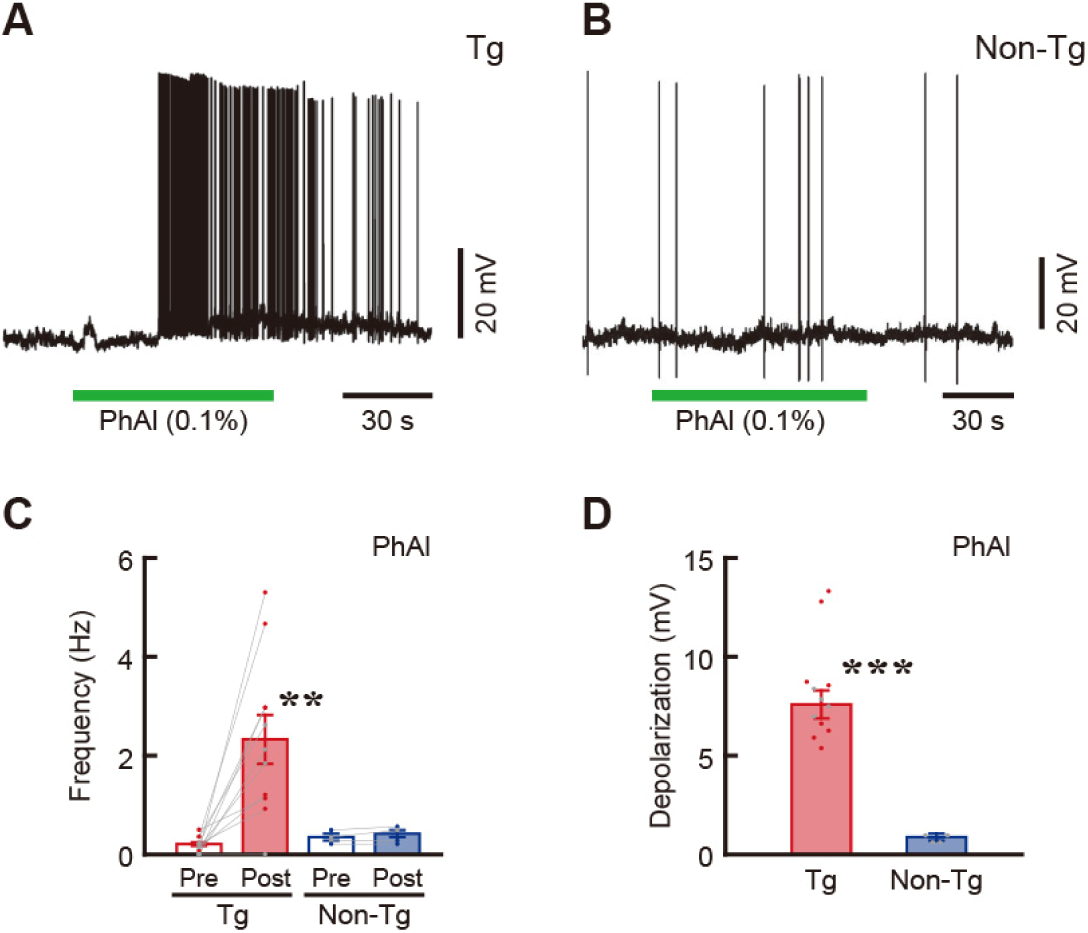
Whole-cell current-clamp recording of a brain slice preparation. (**A**) Excitatory effect of PhAl (0.1%) on the membrane potential of a NE neuron obtained from a Tg mouse. (**B**) Lack of effect of PhAl (0.1%) on the membrane potential of a neuron from a non-Tg mouse. (**C, D**) Bar graphs showing the firing frequency before (pre) and after (post) PhAl application (**C**) and amplitude of PhAl-induced depolarization (**D**) of NE neurons. The firing frequency of neurons in the Tg mice was significantly increased from 0.18 ± 0.05 Hz (pre) to 2.34 ± 0.49 Hz (post) by the PhAl application (n = 11, paired two-tailed t-test, t_10_ *=* 4.462, **p = 0.0012). In the non-Tg mice, the frequency was similar between pre- and post-PhAl applications (0.33 ± 0.06 and 0.41 ± 0.09, respectively; n = 4, paired two-tailed t-test, t_3_ = 1.999, p = 0.1395). The amplitude of PhAl-induced depolarization (7.57 ± 0.72 mV) was significantly higher than the non-Tg value (0.89 ± 0.10 mV) (unpaired two-tailed t-test, t_14_ = 5.201, ***p < 0.0005). Data are presented as mean ± SEM.

**Figure 1 - figure supplement 3.**
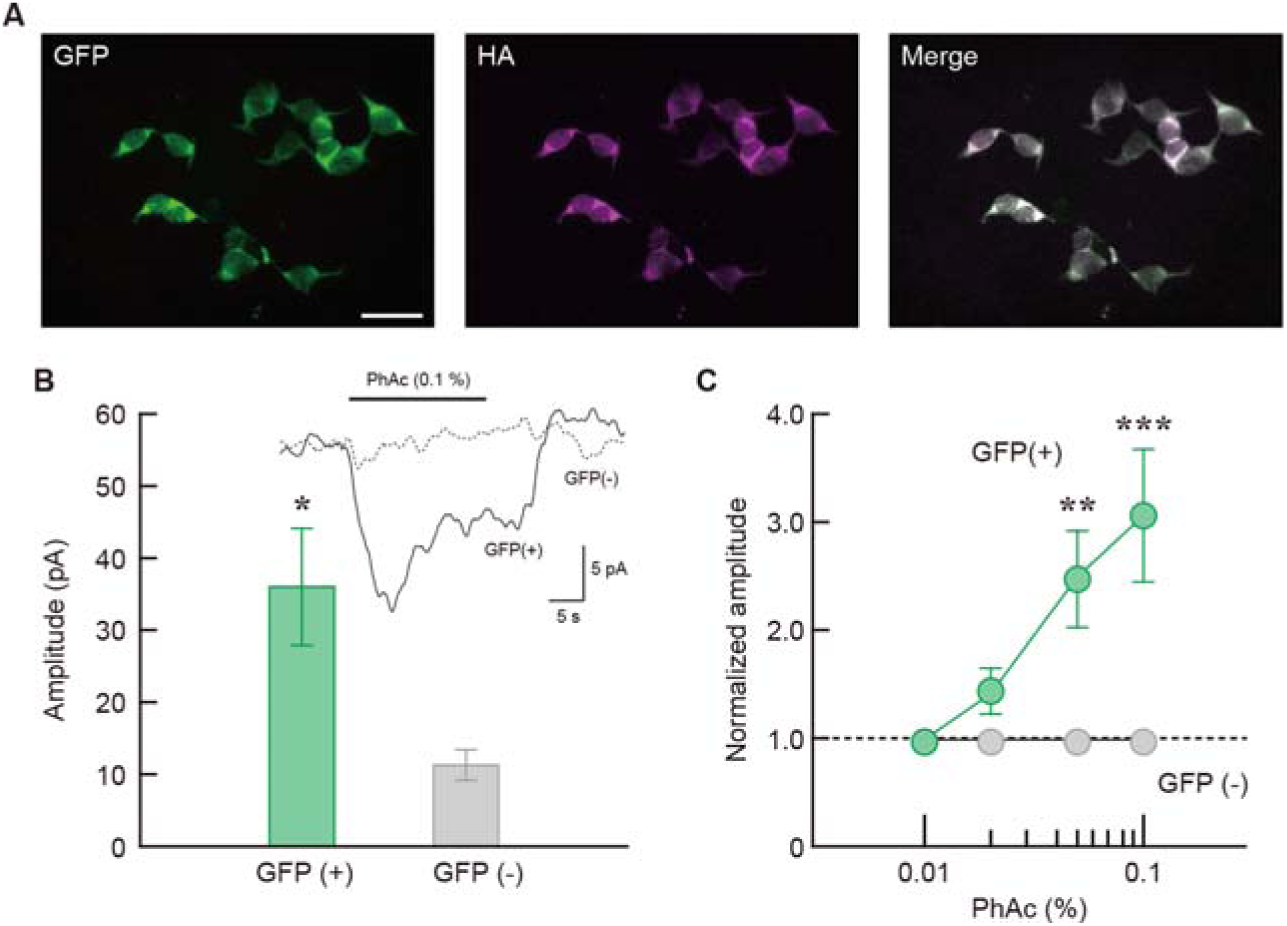
Electrophysiological analysis in cultured cells. (**A**) Expression of IR84a/IR8a complex in HEK293T cells transduced by a lentiviral vector encoding transgene. Immunohistochemistry for EGFP (green) and HA tag (magenta) detected expression of the 2 receptor subunits. Scale bar: 50 μm. (**B**) Ligand-induced inward current of GFP-positive cells expressing the receptor complex. Bar graph showing the current amplitude indicates a significant increase of the amplitude in GFP-positive cells over that of GFP-negative control cells. The mean amplitudes of GFP-positive and GFP-negative cells were 35.77 ± 7.91 pA (n = 11) and 11.14 ± 2.19 pA (n = 14), respectively (Mann-Whitney’s U = 28.00, p = 0.0073). Inset shows mean current traces of the PhAc-induced inward current waveform. Solid and dotted lines indicate the waveform of GFP-positive and negative cells, respectively. (**C**) Ligand dose responses of the GFP-positive (n = 12) and GFP-negative (n = 8) cells. The amplitudes of the ligand-induced currents were normalized in each cell to the amplitude at 0.01% PhAc. PhAc-induced current responses were specific to GFP-positive cells and displayed dose dependency (two-way mixed-design ANOVA, group effect: F_(1, 18)_ = 8.457, p = 0.0094, dose effect: F_(3, 54)_ = 4.396, p = 0.0077, interaction: F_(3, 54)_ = 8.509, p = 0.0001, simple-main effect of dose was significant for GFP-positive cells, F_(3, 54)_ = 12.10, p < 0.0001, but not for GFP-negative cells, F_(3, 54)_ = 0.809, p = 0.4942). The responses of GFP-positive cells at 0.05% and 0.1% PhAc were significantly higher than those of GFP-negative cells (simple-main effect tests of group, F_(1, 72)_ = 7.863, **p = 0.0065, F_(1, 72)_ = 25.09, ***p < 0.0001, respectively). Data are presented as mean ± SEM.

**Figure 2 - figure supplement 1.**
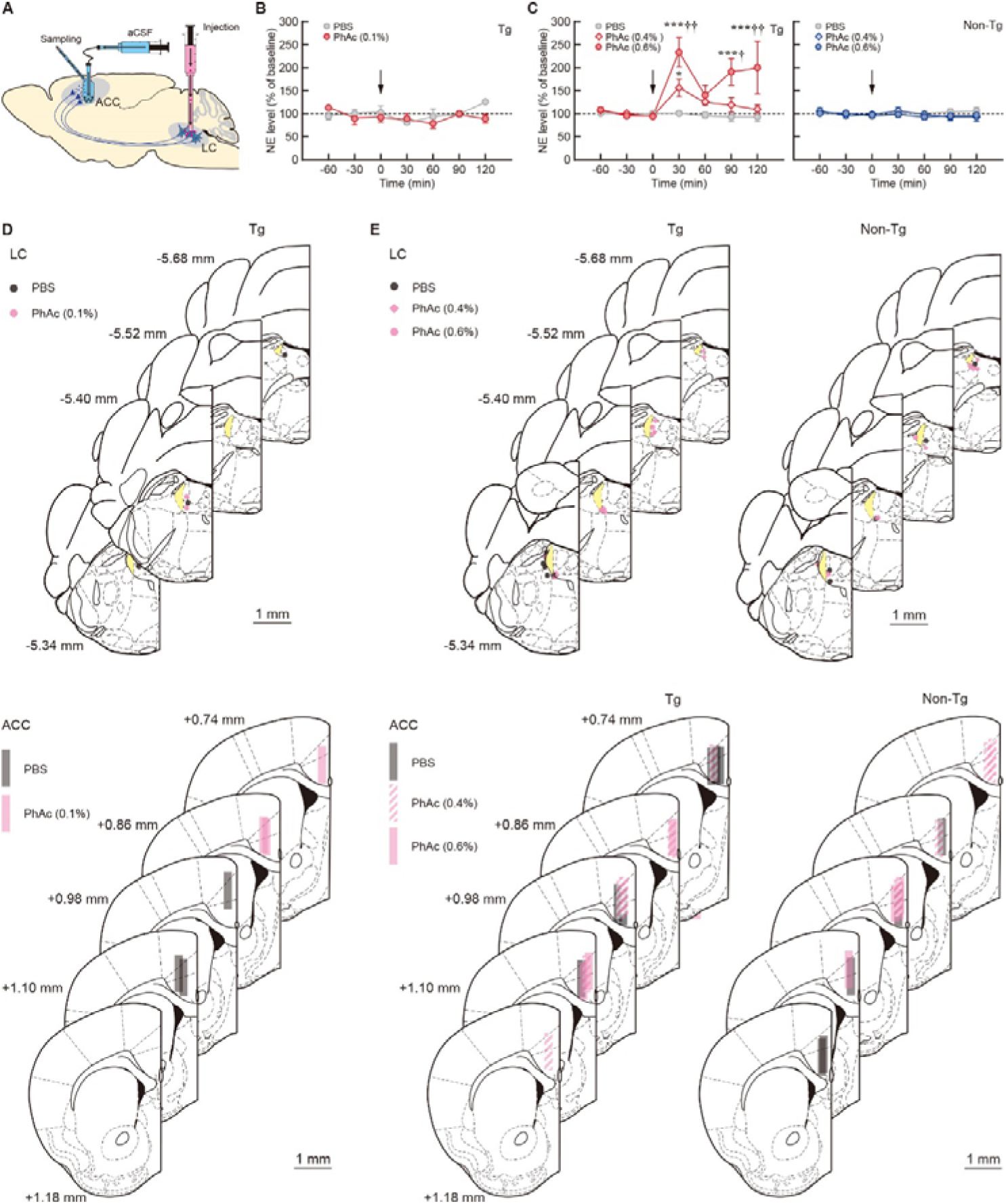
Microdialysis analysis of NE release after PhAc stimulation. (**A**) Diagram for the microdialysis for measuring NE release in the ACC in response to LC activation. (**B**) PhAc solution (0.1%) or PBS was injected into the LC of the Tg mice, and dialysis samples were collected from the ACC. NE levels are expressed as a percentage of each animal’s average baseline levels. LC injection of 0.1% PhAc did not cause a significant increase in NE release in the ACC of Tg mice (n = 4 for each group, two-way ANOVA, drug effect: F_(1, 4)_ = 1.508, p = 0.2868, fraction effect: F_(6, 24)_ = 1.481, p = 0.2268, interaction: F_(6, 24)_ = 1.747, p = 0.1531). (**C**) PhAc solution (0.4/0.6%) or PBS was injected into the LC, and dialysis samples were collected from the ACC. There was no significant difference in average tonic NE concentration (pg/sample) between the two kinds of mice during baseline fractions before the microinjection: Tg, 0.24 ± 0.07 (n = 18), non-Tg, 0.30 ± 0.08 (n = 10) (unpaired two-tailed t-test, t_26_ = 0.518, p = 0.6091). Injection of both 0.4% and 0.6% PhAc into the LC caused a rapid and transient increase in the extracellular NE level in the Tg mice (n = 6 for each group, two-way mixed-design ANOVA, drug effect: F_(2, 15)_ = 9.607, p = 0.0021, fraction effect: F_(6, 90)_ = 6.446, p < 0.0001, interaction: F_(12, 90)_ = 3.269, p = 0.0006). The NE level at the 30-min fraction was significantly increased to approximately 157% and 233% of the baseline level for 0.4% and 0.6% PhAc injection, respectively (t_105_ *=* 2.215, *p = 0.0289 for 0.4% vs PBS, ***p < 0.0001 for 0.6% vs PBS, Holm-Bonferroni test), and magnitude of the effect was dose dependent (^††^p = 0.0059 for 0.6% vs 0.4%, Holm-Bonferroni test). Injection of 0.6% PhAc induced a subsequent elevation in the NE level at a timing delayed after the first peak for 90-min (Holm-Bonferroni test, vs. PBS condition, t_105_ *=* 3.892, ***p = 0.0005; vs. 0.4% PhAc condition, t_105_ = 2.837, ^†^p = 0.0109) and 120-min fractions (vs. PBS condition, t_105_ *=* 4.223, ***p < 0.0005; vs 0.4% PhAc condition, t_105_ = 3.564, ^††^p = 0.0011), suggesting the presence of other complex mechanisms for IR-dependent neuronal activation in addition to the influx of monovalent cations. In non-Tg animals, injection of 0.4% and 0.6% PhAc into the LC did not generate any significant changes in NE release (n = 5 for each group, two-way mixed-design ANOVA, drug effect: F_(2, 12)_ = 0.911, p = 0.4282, fraction effect: F_(6, 72)_ = 0.474, p = 0.8255, interaction: F_(12, 72)_ = 0.359, p = 0.9733). Data are presented as mean ± SEM. (**D**) Placement sites of injection needles and dialysis probes for the experiments with 0.1% PhAc solution. (**E**) Placement sites of injection needles and dialysis probes for the experiments with 0.4/0.6% PhAc solution. Scale bars: 1 mm.

**Figure 2 - figure supplement 2.**
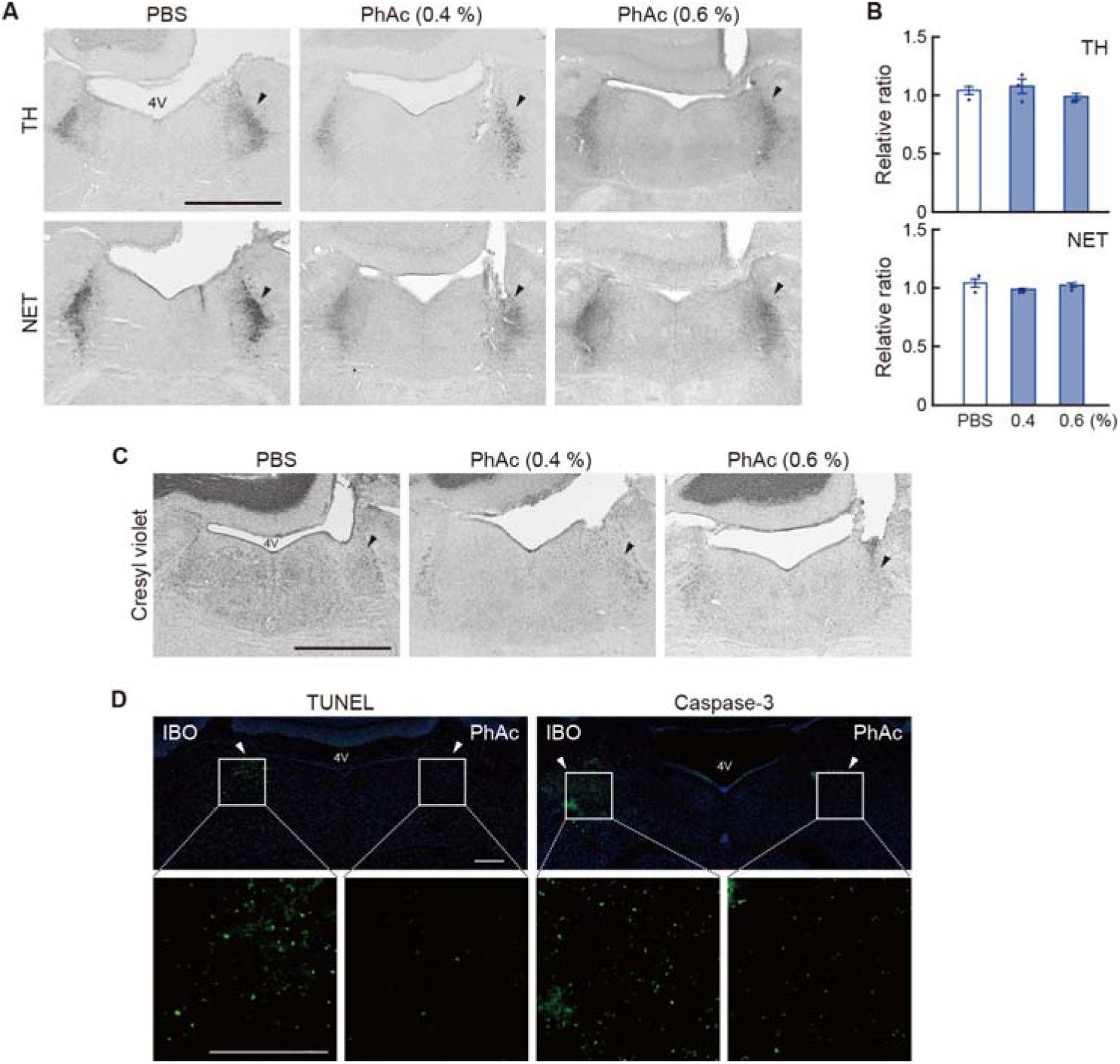
Morphological analysis of LC neurons after PhAc stimulation. (**A**) Morphology of LC neurons in the Tg mice after PhAc stimulation stained by TH/NET. Sections through the LC were prepared from the Tg mice used for microdialysis analysis 7 days after unilateral treatment with PBS or PhAc (0.4/0.6%) and stained for TH and NET immunohistochemistry. LC cells in the PhAc-injected side appear to be normally localized. (**B**) Cell counts for TH and NET-immunopositive cells. The ratio of the number of cells stained for TH or NET in the treated side relative to the intact side was calculated. There was no significant difference in the relative ratios of cell number among the treated groups (n = 3 for each group, one-way ANOVA, F_(2, 6)_ = 1.085, p = 0.3961 for TH and F_(2, 6)_ = 1.426, p = 0.3115 for NET). Data are presented as mean ± SEM. (**C**) Staining of LC sections with cresyl violet, showing no cell damage against LC neurons in the PhAc-treated side. (**D**) Staining for cell death markers. Ibotenic acid (IBO, 1 mg/ml) or PhAc (0.6%) was injected into the LC (0.2 μl/site) of the Tg mice. IBO was used as positive control to detect cell death signals. LC sections were stained with terminal deoxynucleotidyl transferase-mediated dUTP-biotin nick end labeling (TUNEL) or immunostaining for activated caspase-3. The LC areas in the upper images were 4-fold magnified in the lower images. The number of signals in the PhAc-treated side was profoundly lower than in the IBO-treated side, confirming the lack of cytotoxicity of PhAc treatment against LC cells expressing the IRs. Arrowheads indicate the injection sites into the LC. Scale bar: 1 mm.

**Figure 2 - figure supplement 3.**
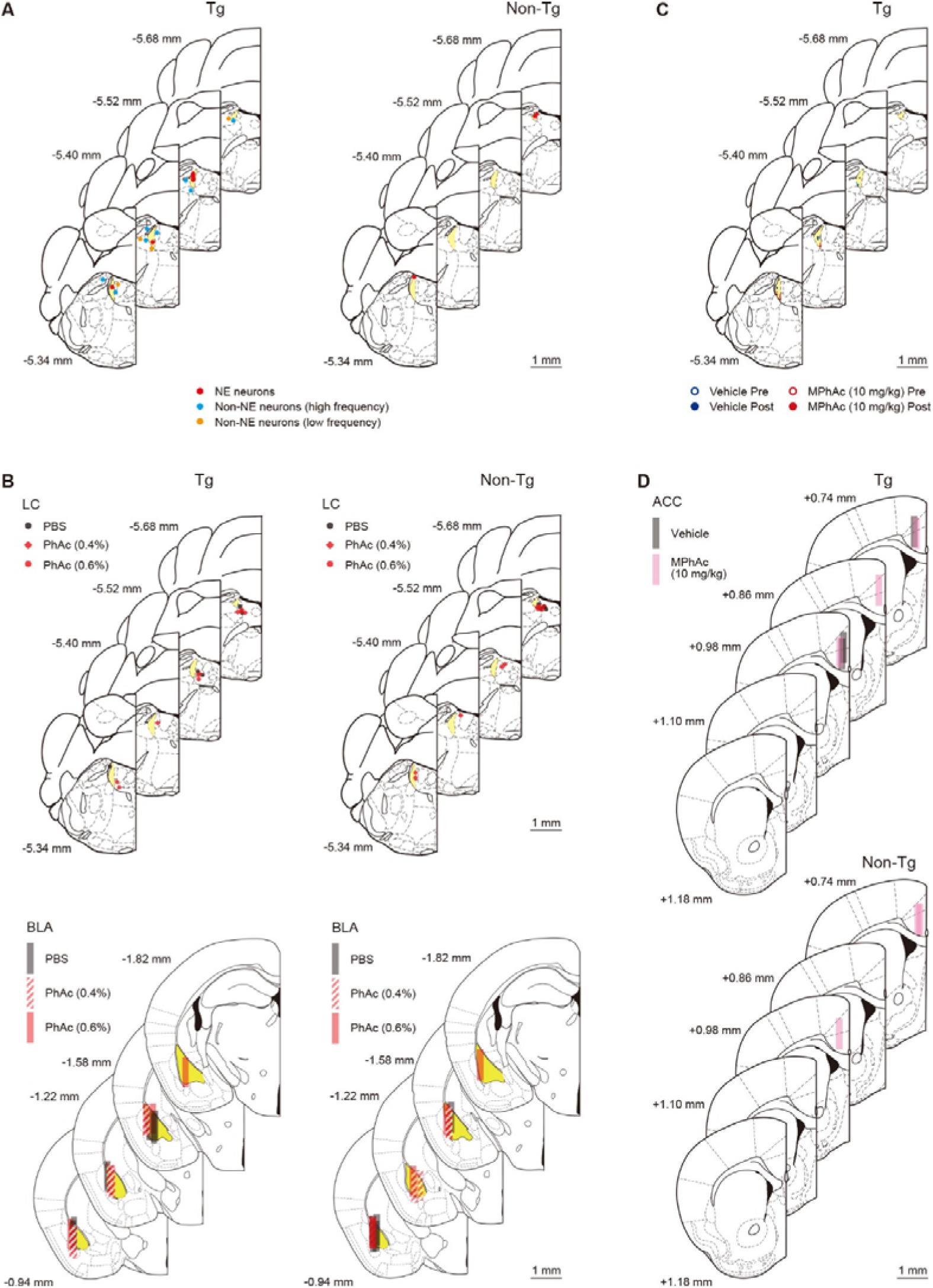
Placement sites of recording electrodes for *in vivo* electrophysiology and injection needles and dialysis probes for microdialysis analysis. (A) Sites of the recording electrodes in the LC for pneumonic injection of PhAc, related to Figures 2B-D. (**B**) Sites of injection needles into the LC and for dialysis probes into the BLA, related to Figure 2F. (**C**) Sites of the electrodes before and after systemic administration of MPhAc or vehicle, related to Figure 2H. (**D**) Sites of dialysis probes after systemic administration of MPhAc or vehicle, related to Figure 2I. The recording sites were marked by pontamine sky blue, and after the experiments the brain sections were prepared and stained by cresyl violet. Scale bars: 1 mm.

**Figure 3 - figure supplement 1.**
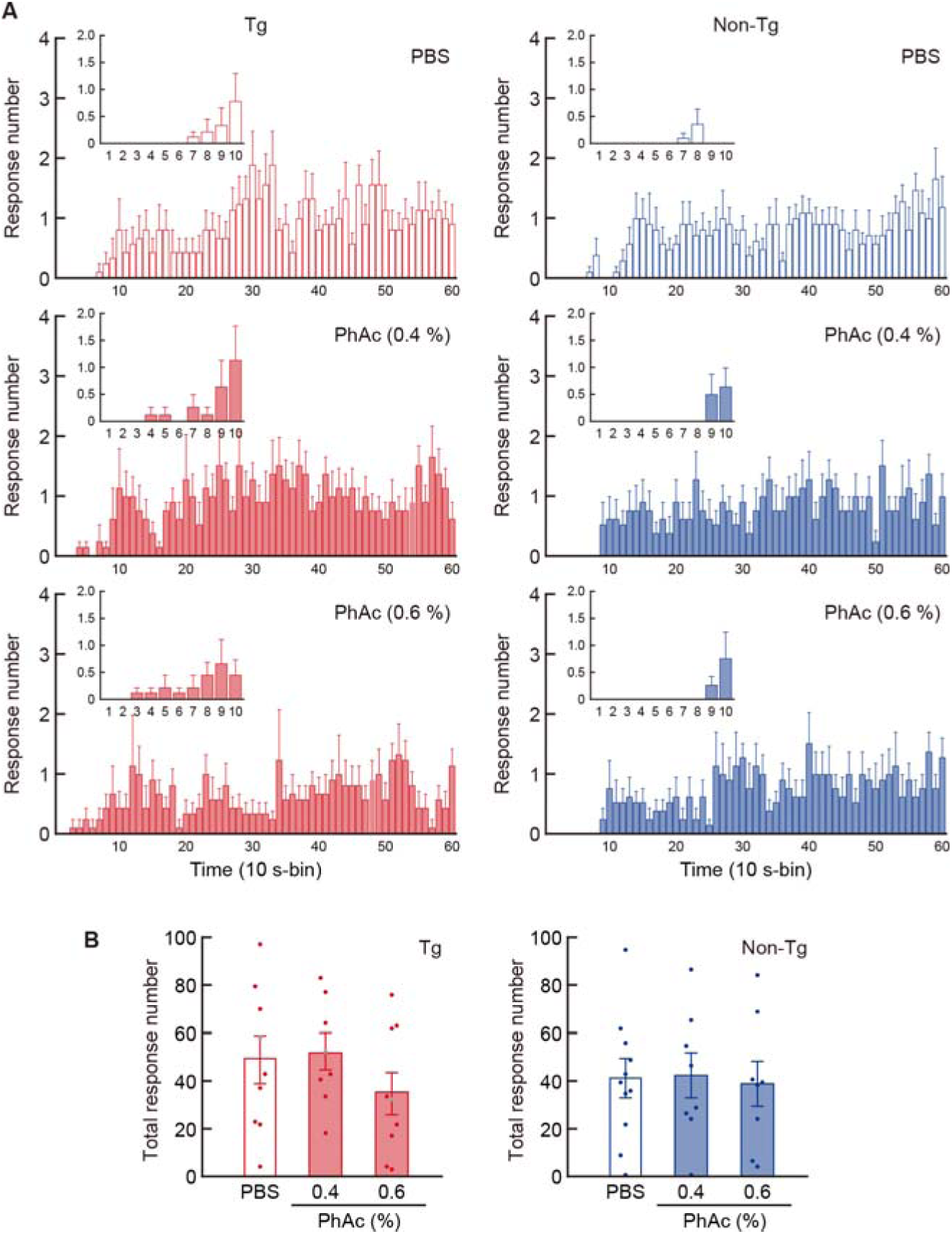
Quantification of aversive responses in the taste reactivity Test. (**A**) Time course of aversive response number. Mice were given intra-LC microinjection of PBS, 0.4% or 0.6% PhAc, and the taste reactivity test was conducted. Behavior was recorded using a digital video camera, and rejection responses (including gaping, chin rubbing, forelimb flailing, paw wiping, and CS dropping) during the 10-min test period were counted. The behavioral data used in Figure 3C was analyzed. The number of responses at a 10-s bin was divided by the number of animals used in each group. Insets show the response number during the early period (< 100 s). (B) Total number of aversive responses during the 10-min test. The number was not significantly different among the three treatment groups in the Tg or non-Tg mice (one-way ANOVA; n = 8-9, F_(2, 23)_ = 0.969, p = 0.3945 for Tg mice; n = 8-11, F_(2, 24)_ = 0.033, p = 0.9677 for non-Tg mice). Data are presented as mean ± SEM. The aversive responses in the Tg mice were observed in the earlier bins in the 0.4% or 0.6% PhAc-injected group compared to the PBS-injected group, although these changes were not seen in the non-Tg mice. The data support the shortening of latency for the initiation of rejection behaviors after 0.4% or 0.6% PhAc treatment in the Tg mice. The similarity of total number of aversive responses during the test period suggests that PhAc treatment before the test does not alter the storage of taste memory in Tg mice.

**Figure 3 - figure supplement 2.**
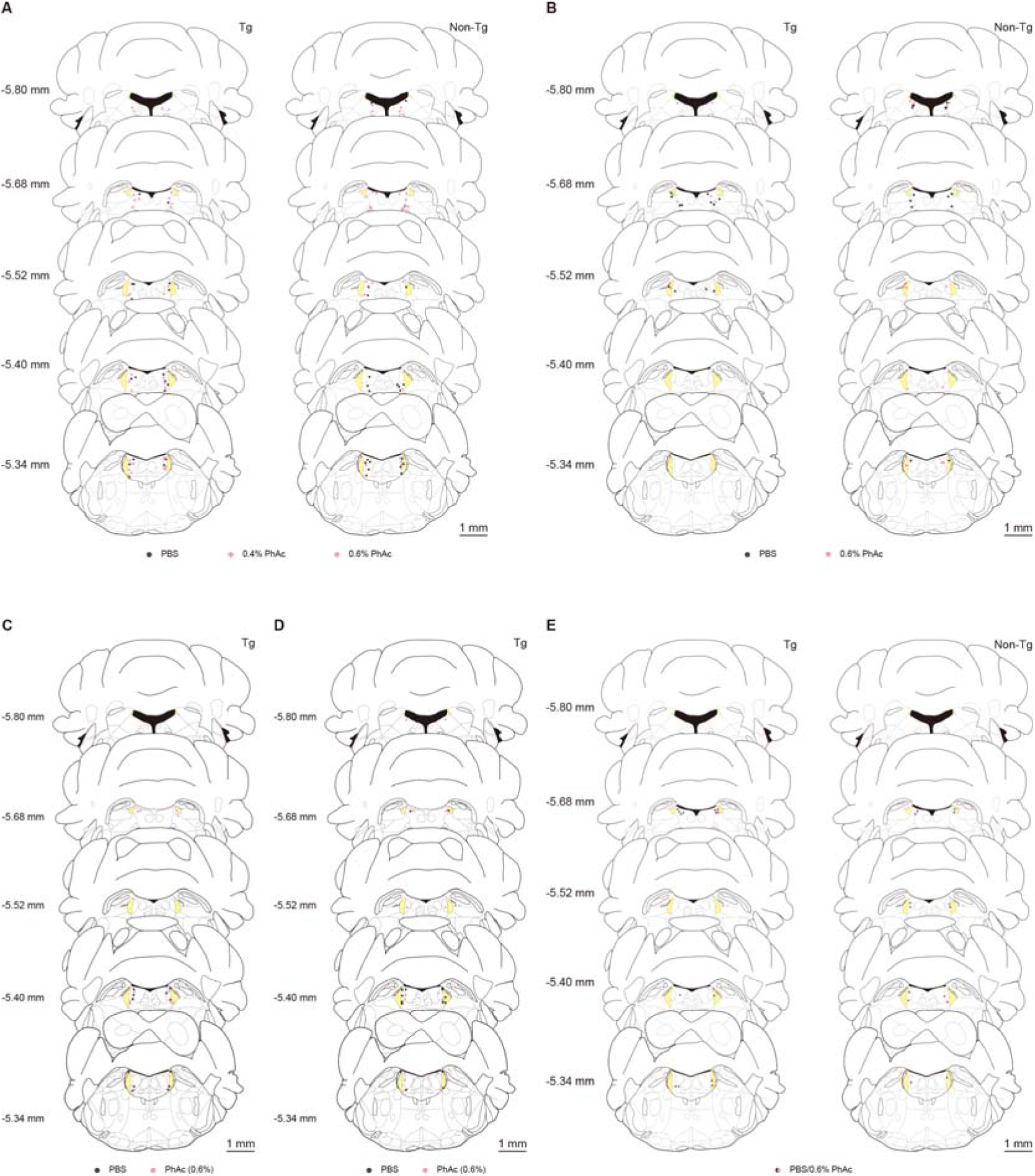
Placement sites of injection needles for behavioral analysis of mice after LC microinjection. (**A**) Sites for the taste reactivity test, related to Figure 3C. (**B**) Sites for the taste sensitivity test, related to Figure 3D. (**C**) Sites for the taste reactivity test of unconditioned mice infused with 0.5 M sucrose, related to Figure 3E. (**D**) Sites for the taste reactivity test of unconditioned mice infused with 0.2 mM quinine, related to Figure 3F. (**E**) Sites for the locomotor activity test, related to Figure 3G. After the behavioral tests, brain sections were prepared and stained by cresyl violet. Scale bars: 1 mm.

**Figure 4 - figure supplement 1.**
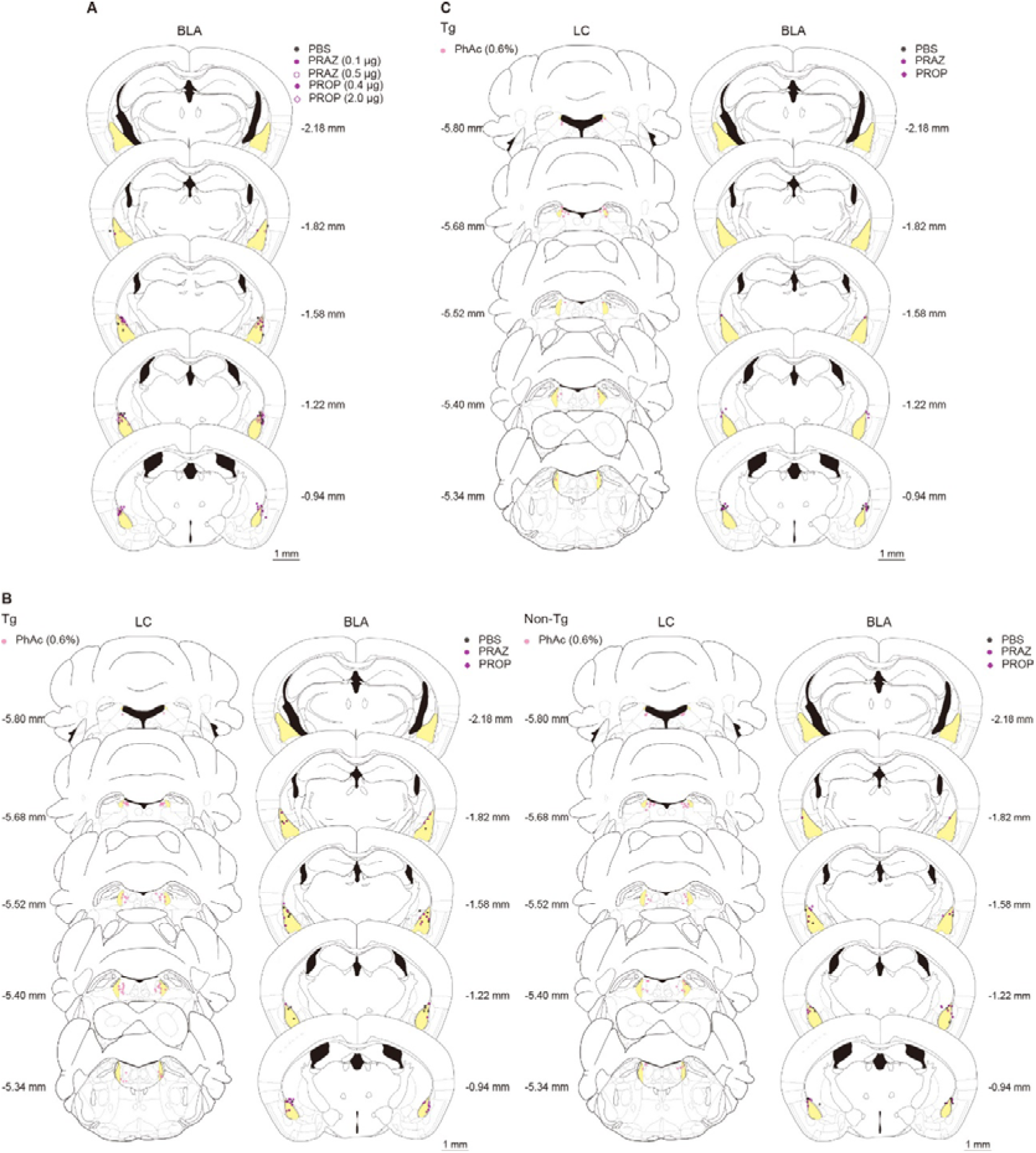
Placement sites of the needles for pharmacological blocking experiments against ligand-induced LC activation. (**A**) Sites for the taste reactivity test in mice that received the treatment of adrenergic receptor antagonists into the amygdala (AMY), related to Figure 4A. (**B**) Sites for the reactivity test after the intra-amygdala treatment followed by LC stimulation, related to Figure 4B. (**C**) Sites for the locomotion test after the intra-BLA treatment followed by LC stimulation, related to Figure 4C. After the behavioral tests, brain sections were prepared and stained by cresyl violet. Scale bars: 1 mm.

**Figure 5 - figure supplement 1.**
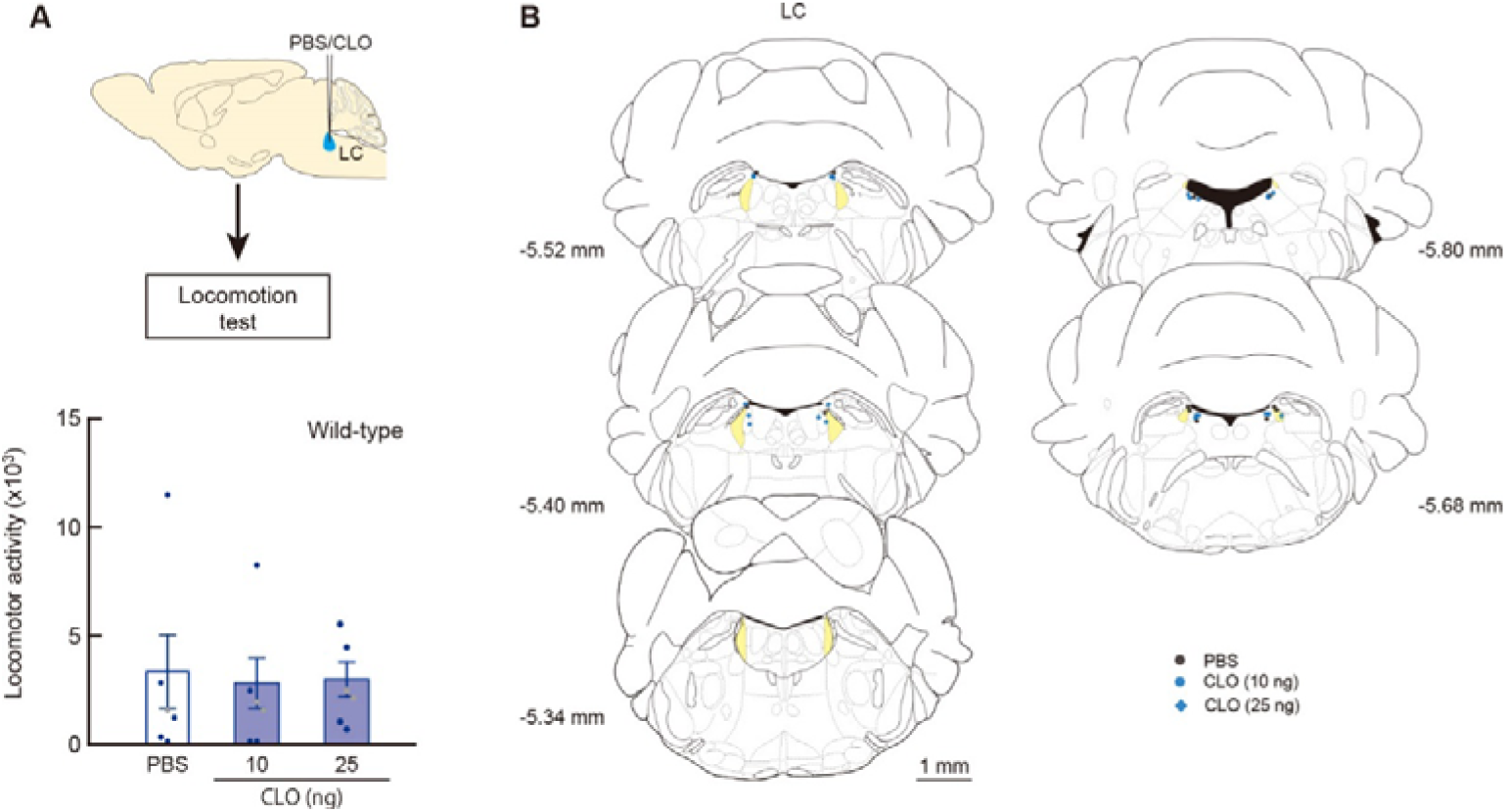
Locomotion test after the intra-LC injection of CLO. (**A**) Locomotor activity. The total number of beam breaks during a 60-min test period was calculated as locomotor activity in response to drug treatment (blocks 1–6) after habituation for 60 min. One-way ANOVA indicated no significant differences among the drug treatments (n = 6 for each group, F_(2, 15)_ = 0.042, p = 0.959). Data are presented as mean ± SEM. Individual data points are overlaid. (**B**) Sites for the locomotion test after drug injection. After the behavioral tests, brain sections were prepared and stained by cresyl violet. Scale bars: 1 mm.

**Figure 5 - figure supplement 2.**
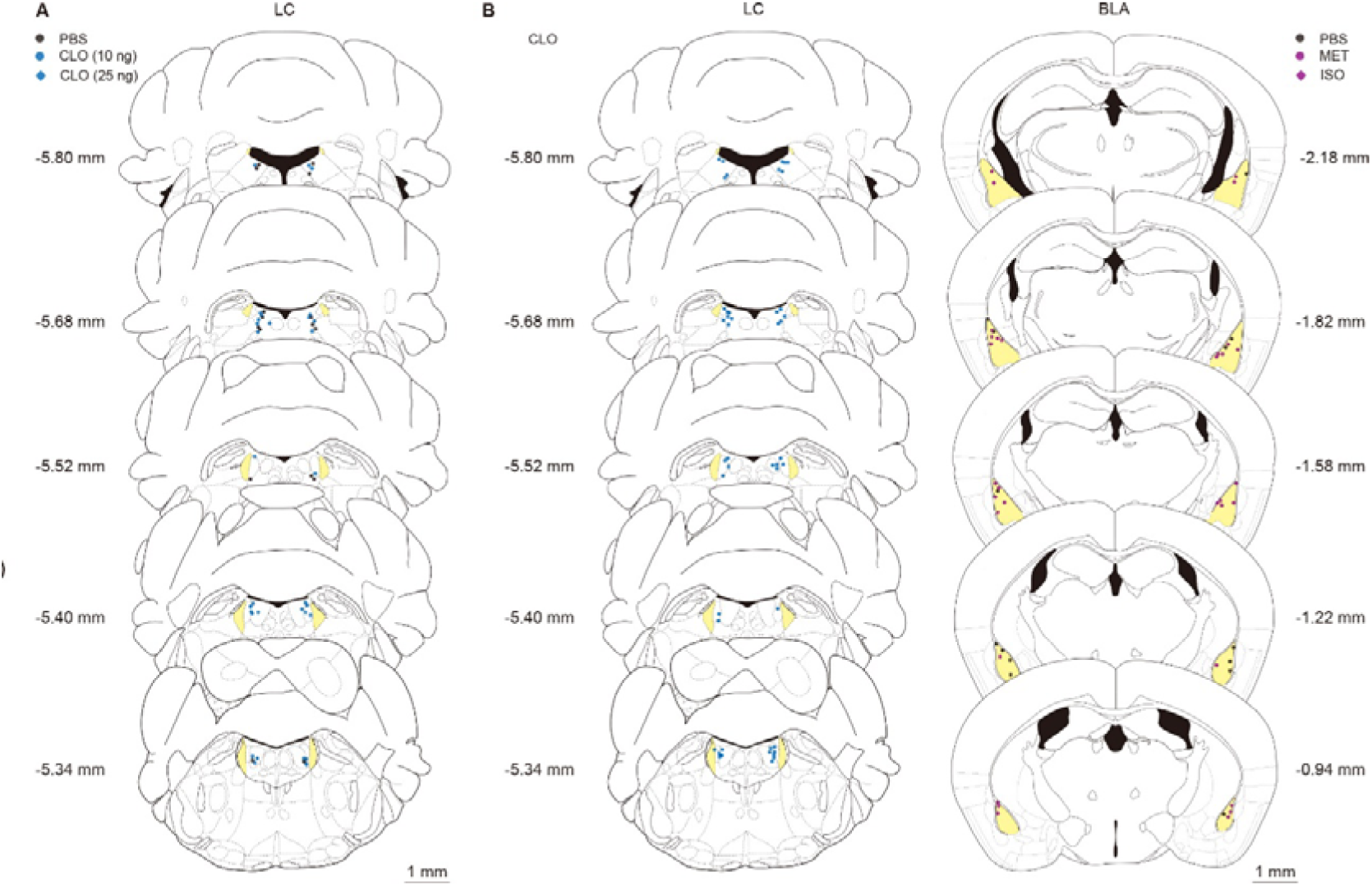
Placement sites of the needles for behavioral tests of mice that received LC pharmacological inhibition. (**A**) Sites for the taste reactivity test after LC injection of CLO, related to Figure 5A. (**B**) Sites for the reactivity test after the LC injection followed by the intra-amygdala treatment of adrenergic receptor agonists, related to Figure 5B. After the behavioral tests, brain sections were prepared and stained by cresyl violet. Scale bars: 1 mm.

